# A single cell atlas of mouse podocytes upon injury identifies kidney zone-dependent responses

**DOI:** 10.64898/2026.02.26.708349

**Authors:** Jeffrey W Pippin, Courtney R. Armour, Diana G. Eng, Uyen Tran, Robert Allen Schweickart, Natalya Kavarina, Kim Dill-McFarland, Oliver Wessely, Stuart J Shankland

## Abstract

There are regional differences in the kidney in focal segmental glomerulosclerosis (FSGS) where podocyte injury is more severe in the juxta-medulla (JM) compared to the outer cortex (OC). Single nuclear RNA-sequencing was performed to determine any regional transcriptomic differences. 1055 differentially expressed genes (DEGs) were identified between healthy OC and JM podocytes. Of 53 podocyte canonical genes, only *Magi1* and *Mapt,* and *Npnt* were higher in healthy OC and JM podocytes respectively. Hallmark pathway analysis showed that in normal mice, healthy JM podocytes are enriched for oxidative phosphorylation, glycolysis and fatty acid metabolism compared to OC podocytes. At day 7 in an experimental FSGS model induced in mice with a cytopathic anti-podocyte antibody, 226 and 225 DEGs were higher in OC and JM podocytes respectively, and 166 overlapped. Five podocyte subclusters were identified, of which the most severe (subcluster 4) was enriched for a senescence phenotype, including the p53 pathway. Inducing FSGS in mice in which p53 was deleted specifically in podocytes had lower glomerular injury compared to diseased wildtype mice. These results are consistent with differences in podocytes in the OC and JM, which might underlie regional differences in FSGS.

## INTRODUCTION

Nephrogenesis occurs in a radial fashion, where it starts in the inner cortex and progresses outward towards the subcapsular cortical region.^1^ Consequently, the earliest glomeruli are formed deeper in the juxtamedullary region (JM) and the younger ones in the cortex (OC). In healthy adult human kidneys, juxtamedullary (JM) glomeruli make up 10-15% of all glomeruli,^2^ with the number being closer to 20% in adult rodent kidneys.^3^ There are several differences between OC and JM glomeruli, with some of them being species-specific. Data from several human populations indicates that the microanatomy of the healthy kidney varies considerably with the mean glomerular volume varying up to 7-fold and the volumes of individual glomeruli within a single kidney varying as much as 8-fold.^4^ For example, human anatomy textbooks^5, 6^ and primary literature^7^ describe that JM glomeruli are 50% or more larger in size than superficial glomeruli. Conversely, Newbold *et al.* reported that in healthy adults, normal JM glomeruli are no larger than the cortical glomeruli.^8^ Similarly, Samuel *et al.* found no differences in size between healthy OC and JM glomeruli in young males, but in advanced age (51-69 years) glomerular volume in the superficial cortex was 20% larger than in the JM when analyzing non-sclerosed glomeruli.^9^ Interestingly, they also reported that this was impacted by race, as JM gloms were larger in blacks compared to whites.^9^ Finally, in living kidney donors, Denic and Rule found that the glomerular volume was largest in the mid cortex (0.0031 μm^3^) compared to superficial cortex (0.0025 μm^3^) and the deep cortex (0.0028 μm^3^) in non-sclerotic glomeruli.^10^ Animal studies in mice, rats, hamsters and dogs reached similar conclusions, where JM glomeruli are reported to be significantly larger than cortical glomeruli and males having larger glomeruli than females.^3, 11^

In addition to size, there are also differences in podocyte number by kidney zone. In non-diseased humans, JM glomeruli have 600-800 podocytes per glomerulus compared to 400-600 podocytes in cortical glomeruli.^4, 12, 13^ Similarly, we reported a higher podocyte density of 404 podocytes/µm^3^ in JM glomeruli compared with 226 podocytes/µm^3^ in OC glomeruli in young adult mice.^14^

Another difference lies in glomerular function. The single nephron glomerular filtration rate (snGFR) is correlated with glomerular volume, where larger glomeruli have higher snGFR compared to smaller glomeruli.^10^ Micropuncture studies in rodents report that the snGFR is higher in the JM glomeruli compared to the OC.^15–17^ Because JM nephrons in animals are larger and have a higher blood flow, some suggest that they contribute to a larger fraction of the total GFR, up to ∼35%.^18^ It remains unclear if this holds in humans, as current methodologies cannot distinguish snGFR between OC and JM glomeruli. But one can speculate that if 85% of glomeruli are in the cortex and have one unit of snGFR and the 15% of glomeruli are in the JM and filter two units of snGFR, then the total is 85*1 + 15*2 = 115 units, the JM glomeruli will contribute about 26% of the total GFR.

Finally, kidney disease can differentially impact OC *vs.* JM glomeruli. The precise single nephron GFR differences between healthy OC and JM human glomeruli is unknown. But the overall GFR is proportional to their fractional nephron number with JM glomeruli contributing ∼10-15% of GFR in healthy glomeruli.^2^ Yet diseases differentially effect kidney zones. Clinical FSGS or hypertension preferentially involves JM glomeruli.^19–21^ This has been confirmed in animal models. We reported in our podocyte depletion model of experimental FSGS that podocyte density decreased by 37% and 45% in OC and JM glomeruli, respectively.^14^

In mice with the Adriamycin model of experimental FSGS, 81.3% of the JM nephrons were injured versus 39.6% of OC nephrons.^22^ Additionally, there were differences in podocyte replacement following podocyte loss in this study, where CXCL12 therapy resulted in a more robust regeneration of OC than JM glomeruli.^22^ Similar regional differences have been reported in additional studies of Adriamycin nephropathy in mice,^23,24^ and rats, ^22^ as well as rats with type 2 diabetic kidney disease^25^ and in spontaneously hypertensive rats.^26^ Finally, following a reduction in nephron number in rats, the snGFR was higher in JM glomeruli compared to cortical glomeruli.^16^

Several causal hypotheses have been proposed to explain the regional JM and OC glomerular differences including differences in hemodynamics, oxygen tension, anatomic and functional loads, and size.^8, 27, 28^ However, a major knowledge gap remains in understanding if these differences are paralleled by differences in gene expression between OC and JM podocytes in normal kidneys and during disease. To address this, we performed single nuclear RNA-sequencing (snRNA-seq) of kidneys from healthy mice and mice with experimental FSGS with podocyte depletion by separating the kidneys into JM and OC regions.

## METHODS

### Animals

Mice were bred and socially housed in the animal care facility at the University of Washington under specific-pathogen-free conditions with *ad libitum* food and water. All animal protocols and procedures were approved by the University of Washington Institutional Animal Care and Use Committee under protocol 2968-04.

### Model of podocyte injury for snRNAseq

For the single-nuclear RNA-sequencing and immunostaining studies, four-month-old male and female mixed background mice (n = 12) were randomly assigned to experimental groups of normal (n = 4), D7 of FSGS (n = 4), and D28 of FSGS (n =4), with 2 male and 2 female in each cohort. Induction of FSGS was performed by administering 2 consecutive doses of cytopathic sheep anti-podocyte antibody, 24 hours apart, at 7 mg/20 g body weight.^14, 29, 30^ Individually pooled spot urines were collected for each animal prior to induction of disease and on days 7, 14, and 28 prior to euthanasia.

### Podocyte-specific ablation of p53 in mice

p53 was specifically deleted from podocytes as follows: inducible podocyte specific p53^fl/fl^ (F1: *Podocin-rtTA|tetO-cre|p53^fl/^*^fl^, n =3 male) and wildtype control (F1: *Podocin-rtTA|tetO-cre|p53^+/+^,* n = 4 male) mice were generated by breeding two parallel strains of mice as shown in Figure 5A. Podocin-rtTA or tetO-cre female mice were initially crossed with p53^fl/fl^ male mice, with resulting offspring bred and selected for a maternal strain of podocin-rtTA|p53^+/+^ (wildtype) or podocin-rtTA|p53 ^fl/fl^ (mutant) mice and a paternal strain of tetO-cre|p53^+/+^ (wildtype) and tetO-cre|p53 ^fl/fl^ (mutant) mice. These parallel strains of mice were then bred to generate F1 experimental offspring. Podocin-rtTA (FVB/N-Tg(NPHS2-rtTA2*M2)1Jbk/J, Stock No: 008202),^31^ TetoCre (B6.Cg-Tg(tetO-cre)1Jaw/J, Stock No: 006234)^32^ and p53fl/fl (B6.129P2-Trp53tm1Brn/J, Stock No: 008462)^33^ mice were all originally purchased from The Jackson Laboratory. Adult 3-month-old experimental animals were all provided ad libitum 625mg/kg doxycycline chow (TD.01306, Envigo) for 3 weeks to induce conditional knockout of p53 in podocytes in mutants. At least one week of clearance on regular chow was provided prior to the induction of FSGS as above at 10 mg/20 g body weight. Urines were collected prior to induction of disease and on D14 prior to euthanasia.

### Podocyte reporter mice for urinary podocytes

To determine if podocyte depletion in this model of injury was accompanied by podocyte detachment into the urine, 3- to 4-month-old male and female mixed background *Nphs1-FLPo|FRT-EGFP* mice as previously described^34, 35^ were utilized for their podocyte specific constitutive expression of EGFP to facilitate podocyte isolation. Induction of FSGS was performed as above at 10 mg/20g body weight on n = 8 animals. Spot urines were collected at 2-hour intervals 4 times per day on days 3 and 4 of FSGS, and twice on day 5 of FSGS, prior to euthanasia. Four animals were euthanized without the induction of FSGS to serve as baseline reference animals.

### Sample processing and snRNA-sequencing

Single-nucleus mRNA sequencing (snRNA-seq) was performed on mouse kidney tissue collected from the three injury time points (t=0, t=7d and t=28d) and micro-dissected into juxtamedullary (JM) or outer cortical (OC) zones of the kidney following the protocol of the KPMP.^36^ Library preparation and sequencing were performed using the 10x Genomics Chromium platform.

### snRNA-seq analysis

Sequencing reads were processed through the default pipeline for Cell Ranger (v6.0.0)^37^ with include-introns=true flag and aligned to the reference mouse genome (version mm10) to generate raw count matrices. Resulting 10x files were processed with scanpy (v1.10.2)^38^ to identify doublets with scrublet (0.2.3),^39^ remove genes found in fewer than 3 cells, remove cells with less than 500 or more than 5000 genes, and remove cells with greater than 10% mitochondrial reads. Scanpy defaults were used unless otherwise specified. A total of 256,126 high quality cells expressing 17,790 genes were identified for downstream analysis. Cells were integrated using scVI (v1.1.5)^40, 41^ with sampleID as a categorical covariate and total counts and percent mitochondrial reads as continuous covariates, then reads were normalized to median total counts and log transformed.

### Cell type annotation

Cells were clustered using UMAP and the Leiden method^42^ where 18 distinct clusters were identified at resolution 0.7. The rank_genes_groups function in scanpy was used to identify marker genes for each cluster. Clusters were manually annotated based on the marker genes and prior mouse kidney cluster identification.^43^ Proportions of cells in clusters were compared between kidney zones (OC, JM) and timepoint after FSGS induction (D0, D7, D28) using an overall Kruskal-Wallis test followed up pairwise Wilcoxon rank sums tests. Significance was defined at p-value < 0.05. One of the 18 clusters did not exhibit strong marker genes and had lower quality metrics; it was classified as low-quality RNA cells and omitted from further analysis.

### Differential gene expression analysis

Due to limited sample size and high sequencing depth, which results in over-powered single-cell analyses, we used a pseudobulk approach to identify differentially expressed genes (DEGs) between kidney zones (OC, JM) and FSGS disease (normal, D7, D28). For each cell type of interest, raw gene counts were aggregated by sample using the Aggregate Expression function in the Seurat R package (v5.4.0). Gene counts were normalized using trimmed mean of *M* values (TMM) normalization, filtered to protein-coding genes with at least 1 count per million in at least 3 samples, and converted to log_2_ counts per million with quality weights using ‘voom’ (v3.66.0).^44^ For podocytes, this resulted in 14,470 genes for analysis from 24 samples that was then used for mixed effects regression with kimma (v1.6.3).^45^ Models compared the six-level concatenated kidney zone plus disease variable (normal JM, D7 JM, etc.) with random effect for mouse to account for JM and OC from the same animal. Prior analyses showed sex differences in kidney gene expression; therefore, we included sex in the model as a covariate. The model per gene was as follows:

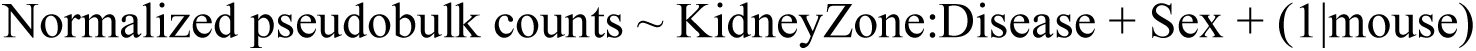

Benjamini-Hochberg p-value correction was applied to the results and DEGs were defined at FDR < 0.05. Pathway enrichment was performed for DEGs of interest using hypergeometric enrichment against the Broad Molecular Signatures Database (MSigDB) Hallmark gene sets.^46^ Pathway significance was defined at FDR < 0.05.

### Podocyte sub-clustering

The 7,306 cells in the podocyte cluster were subset, re-integrated with scVI, and clustered using UMAP with the Leiden method to create 5 podocyte subclusters at resolution 0.35. Using the scanpy rank_genes_groups function to identify marker genes, we identified a large subset of healthy podocyte cells (n=4307, 60%), a subset of PEC-like cells (n=1491, 20.4%), and a subset of injured podocytes (n=398, 5.4%). Marker genes comparing the healthy and injured podocyte clusters were identified, and hypergeometric enrichment was conducted on these marker genes as described for DEGs.

### Urinary podocytes for liquid biopsy

Spot urines were collected from individual mice directly into cryotubes with light depression of the bladder while scruffed. Within 15 minutes of collection, each two-hour sample was spun down at 1000 RPM (300 g) for 5 minutes at 4 °C. The supernatant (urine) was removed from the cryotube and stored at −80C for urine analysis. Remaining cell pellets were resuspended in 50ul CryoStor CS10 solution (#210373, Biolife Solutions, Bothell, WA) and placed into Cryofreezing chambers at −80 °C to freeze at −1 °C/minute. Cryovials were then transferred to liquid nitrogen for storage until FACS sorting. Approximately 1-2 ml of total urines were collected from each animal over 10 cumulative spots on days 3-5 of FSGS. Kidneys were collected on FSGS day 5 from each corresponding mouse and were digested into single cell suspensions to obtain resident EGFP-positive podos from the kidney as we have reported ^47–49^ Cryopreserved urinary cells were thawed, and urine collections from days 3-5 for each individual mouse were pooled. Urinary cells and resident kidney cells from kidney digests were separately resuspended in BD FACS^TM^ Pre-Sort Buffer and subjected to FACS analysis on a BD FACS Discover S8 sorter. EGFP-positive podocytes and EGFP-negative urine and kidney cell fractions were collected separately. Sorted urine and kidney pellets containing 50-1,000 cells were resuspended in 7ul PBS in 8 tube strips and frozen at −80°C. Bulk RNA-sequencing was then performed by Psomagen, (Rockville, MD, USA) where indexed cDNA libraries were generated directly from 1–1,000 intact cells from the urine and kidney cell pellets using the SMARTer Stranded RNA-Seq Kit (Takara Bio USA, Inc., San Jose, CA, USA). Next-generation sequencing (NGS) was then performed on the Illumina platform.

### Urine Albumin to Creatine Ratio (ACR)

ACR was calculated as previously described using a radial Immunodiffusion assay for albumin^50^ and the Creatinine (urinary) colorimetric assay kit (#500701, Cayman Chemical).

### Immunostaining

Indirect immunoperoxidase and immunofluorescence staining were performed on 4μm thick formalin fixed paraffin-embedded (FFPE) mouse kidney sections as previously described.^51^ Briefly, paraffin was removed from formalin-fixed sections using Histoclear (#HS-200, National Diagnostics) and rehydrated in a graded series of ethanol. Antigen retrieval was performed by boiling sections in in antigen retrieval buffer (Citric acid buffer pH 6.0 or EDTA buffer pH 6.0, pH 8.0). Sections were blocked for endogenous peroxidase (3% hydrogen peroxide) and Avidin/Biotin Blocking (#SP-2001, Vector Laboratories). Non-specific antibody binding was blocked with Background Buster (#NB306, Accurate Chemical & Scientific). Antibodies were diluted in 1% IgG free BSA in PBS and were incubated overnight at 4°C. Secondary horseradish peroxidase conjugated antibodies and streptavidin conjugates were detected diaminobenzidine (#112090250, DAB, Fisher Scientific). In some cases, Periodic acid Schiff’s (PAS) staining was performed as a counterstain. Slides were dehydrated in ethanol and mounted with Histomount (#HS-103, National Diagnostics, Atlanta, GA).

Immunofluorescence staining was performed on paraffin sections, following deparaffinization and antigen retrieval with citric acid buffer (pH 6.0), samples underwent the same blocking and antibody incubation protocol as described for immunoperoxidase staining. After secondary antibody incubation, the samples were incubated with tetramethylrhodamine-tyramide at a 1:100 dilution (TSA-Vivid kit, (#7523, R&D Systems) at RT for 10 min. All immunofluorescence samples were mounted using Vectashield mounting medium with 4′,6-diamidino-2-phenylindole (DAPI) (#H-1200, Vector Laboratories).

Primary antibodies and analysis of immunostaining.

To identify podocyte injury and determine glomerulosclerosis, p57/PAS staining was performed, the following antibody was applied: rabbit anti-p57 (#sc-56341, 1:1000, Santa Cruz Biotechnology). Nuclear staining for p57 was used to calculate podocyte number in a glomerular cross-section. Podocyte number was calculated by using the “Analyze particles” feature in the Fiji program. In brief, a threshold was set and applied to the images. Then, the glomerular area of each glomerulus was delimited manually, and the total “particles” (corresponding to p57-positive nuclei) were automatically counted as well as the average size of the particles. Data was presented as average number of p57-positive cells per glomerulus. For p57/PAS staining, and all other stains, glomeruli were separated into the outer cortex (OC) and the juxta-medullary (JM) region for quantification purposes. 20 JM and 40 OC glomeruli were counted in each section, and a total sum of OC and JM glomeruli served as a representation of the entire section

To outline OC and JM glomeruli in murine and human kidney samples, staining with NAPSA (#orb519989, 1:25, Byorbit) was performed on paraffin sections as described above. Kidney senescent cells were marked by the following antibodies: rat anti-p16 INK4a-N-terminal (#ab211542, 1:1000, Abcam), rat anti-p53 (#ab241566, 1:1000, Abcam), anti-rabbit serpine-E1 (#NBP1-19773, 1:500, Novus Biologicals) and rat anti-p21 (#ab107099, 1:200, Abcam). Frozen sections were used to measure SA-β-galactosidase activity in kidney, sections were treated in accordance with the manufacturer’s protocol for the SA-β-gal staining kit (#9860, Cell Signaling Technology) as previously described.^52^

Kidney endothelial cells were marked by a rabbit anti-ERG antibody (#ab92513, 1:100, Abcam,). To outline the kidney architecture, biotinylated collagen IV antibody (#1340-08, 1:100, Southern Biotechnology) was applied overnight as second primary antibody. Jones basement membrane staining was performed by the University of Washington Pathology Research Services Laboratory on tissue embedded in paraffin to determine extracellular matrix.

### Statistics

Statistical analysis was performed using GraphPad Prism 10.0 (GraphPad Software, Inc., San Diego, CA). t tests were performed when comparing only two groups of data, whereas repeated-measures one-way ANOVAs were performed when making multiple comparisons. A P value threshold of 0.05 was used to determine statistical significance and data are shown as the means ± SD.

## RESULTS

### Mouse model of FSGS and snRNA-sequencing

To investigate the impact of experimental FSGS at the single cell level, 4-month old young mice of both sexes were administered 2 consecutive doses of cytopathic sheep anti-podocyte antibody as reported previously.^14, 29, 30^ Kidneys from normal healthy mice (t=0) along with kidneys 7 days and 28 days after the second dose were harvested and dissected into separate outer cortex (OC) the juxtamedullary (JM) regions following the protocol of the KPMP^36^ (**Figure 1A**). Staining of the sheep anti-podocyte antibody for sheep IgG deposition confirmed that the antibody targeted podocytes in all glomeruli in both the OC and JM, with no differences in intensity by region (**Figure 1B**). We then performed single nuclear RNA-sequencing (snRNA-seq) following the protocol established by the KPMP consortium.^36^ After quality control filtering and preprocessing at each timepoint, region and sex, ample nuclei were obtained for analysis yielding a total of 256,126 nuclei (**Supplemental Figure S1A,B**). Following data integration and clustering of the nuclei using well defined cluster marker genes established for the kidney (**Table 1**), 18 separate clusters comprising glomerular, tubular, stromal, and immune cell types were captured (**Figures 1C, D**). The regional distribution (OC *vs.* JM) was as expected, with the PT3, PT3_Inj_, TAL and Loop of Henle cells being enriched in the JM (**Figure 1E,F**). Except for the proximal tubule populations, the other cell types were relatively evenly distributed by sex (**Supplemental Figure S1C,E**). With respect to the distribution based on disease progression, the clusters were relatively evenly distributed between t=0, 7d and 24d post injury, except for the proximal tubule populations (**Figure 1G** and **Supplementary Figure 1B,D**). As podocytes were the primary target of the cytopathic sheep anti-podocyte antibody, we first focused on the podocyte cluster.

**Figure 1:**
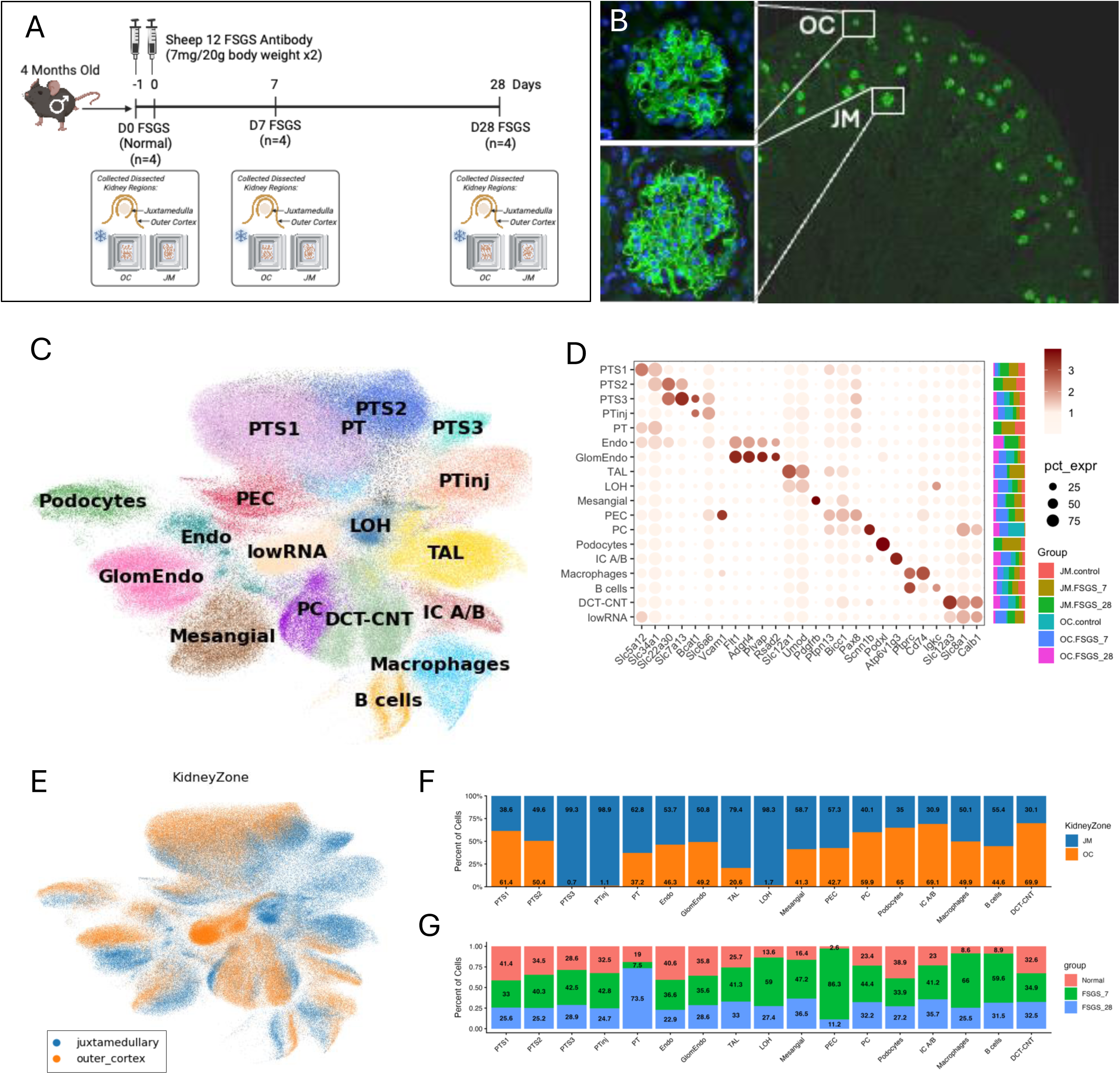
Experimental design and data processing. **(A)** *Experimental Design*. 4-month-old mice were injected with a sheep cytopathic anti-podocyte antibody that induces podocyte injury and loss leading to FSGS. The kidneys of normal mice that did not receive antibody (day 0) and FSGS mice at days 7 and 28 were dissected into separate outer cortical (OC) and juxtamedullary (JM) regions for RNA isolation as shown **(B)** *Sheep IgG staining*. Sheep IgG staining shows that the disease inducing antibody localizes to podocytes in a similar pattern in the OC and JM. **(C)** *Uniform Manifold Approximation and Projection (UMAP)*. The UMAP comprised 256,126 cells after integration, separated into 18 clusters including glomerular, stromal, and immune cell types. **(D)** *Dot plot of marker genes*. The marker genes used to identify individual cell types for the 18 clusters are shown; the size of the dot indicates the proportion of cells in each cluster that express each gene, and the color of the dot represents the mean expression of the gene in each cluster normalized across the clusters. The bar plot on the right represents what proportion of the cells in each cluster from the OC or JM in normal mice and at different FSGS timepoints. (**E**) *Uniform Manifold Approximation and Projection (UMAP)*. The UMAP for regional distribution (OC, orange *vs.* JM, blue) shows even distribution, but with the PT3, PT3_Inj_, TAL and Loop of Henle cells being enriched in the JM region. (**F**) *Regional Distribution of Clusters*. Box plots show the percentage of cells found in each cluster in the OC (orange) and JM (Blue) regions (**G**) *Distribution of Clusters Based on Disease Progression*. Box plots show the percentage of cells found in each cluster at t=0 (pink), FSGS day 7 (green) and FSGS day 28 (blue).

### Differences in Podocyte Gene Expression Between JM and OC in normal mice

Comparison of the regional distribution of the podocytes showed a clear separation between JM and OC in the UMAP with 65% of the podocytes originating from the OC and 35% in the JM (**Figure 1E,F**, **Figure 2A**). As it is currently unknown whether the transcriptome of podocytes in the JM and OC of normal healthy mice differ, we performed a pseudo bulk analysis focusing on the podocytes from normal mice before injury induction (t=0). Regression modeling identified a total of 1,055 differentially expressed genes (DEGs) between the JM and OC in normal mice (**Table 2**) and highlighted in the volcano plot in **Figure 2B** (FDR < 0.05). Novel aspartic proteinase of the pepsin family A is encoded by the *Napsa* gene and the protein stains proximal tubules in a granular cytoplasmic pattern^53^ We identified *Naspa* as a novel marker for cells of the JM region and the gene was highly expressed in podocytes of the JM compared with podocytes of the OC region (**Figure 2C**), which was further validated at the protein level by immunostaining (**Figure 2D**). Moreover, several transporter genes were enriched in specific kidney regions with *Slc22a19*, *Scl12a1*, *Slc22a7*, and *Slc22a13* in the JM and *Slc5a12*, *Slc5a2*, *Scl7a8*, and *Slc7a7* in the OC.

**Figure 2:**
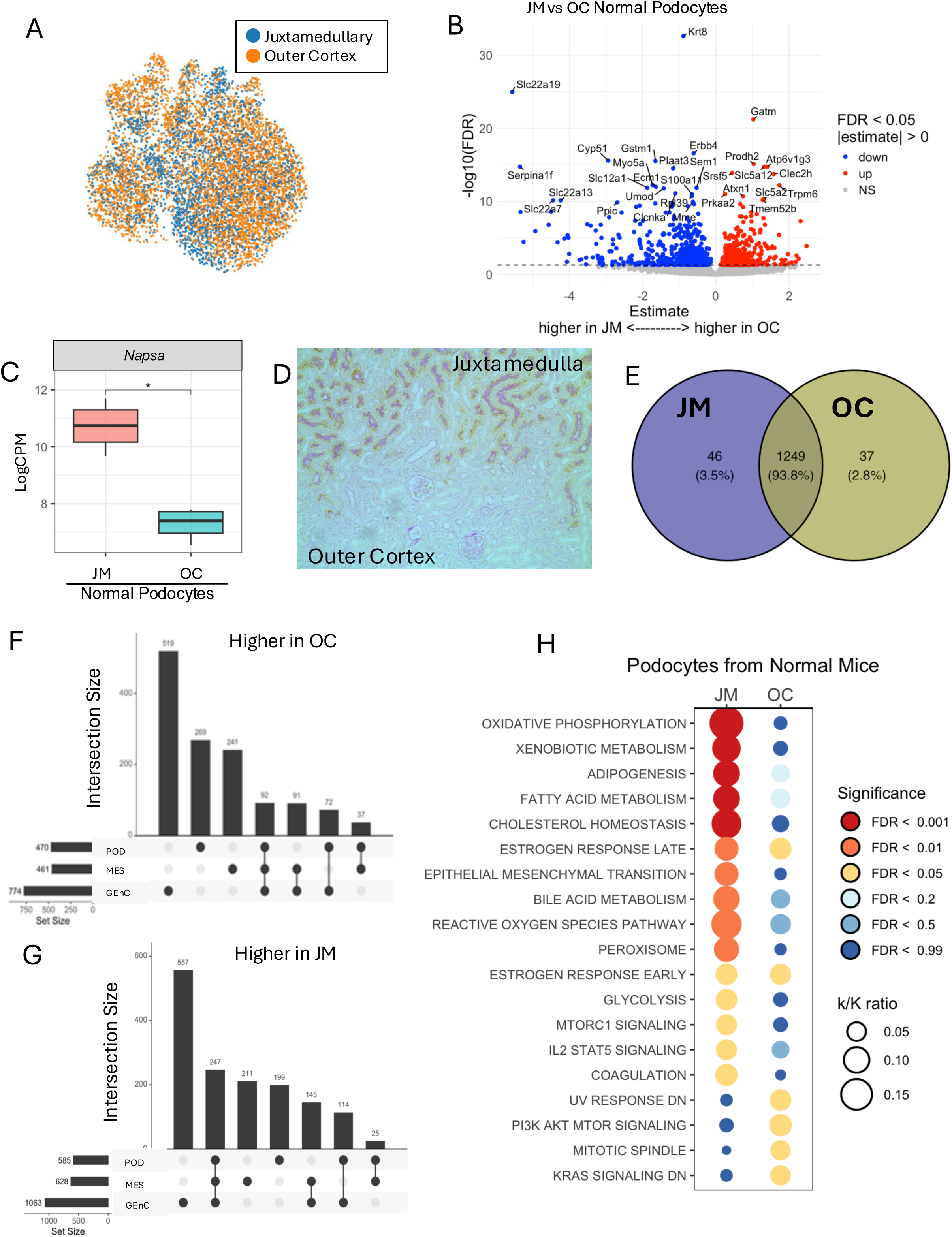
Transcriptional Differences in Podocytes by Kidney Zone in Normal Mice. **(A)** *UMAP*. There was clear separation of nuclei from the juxtamedullary (blue) and cortical (orange) regions. The Venn Diagram shows that 93.8% of DEGs were shared between normal podocytes in the JM and OC, with each zone having 46 and 37 unique genes respectively. (**B**) *Volcano Plot*. A Volcano plot of differentially expressed genes (DEGs) between the JM and OC of normal mice shows a total of 1,055 DEGs (FDR < 0.05). Red indicates genes expressed higher in the OC and blue indicates genes expressed higher in JM. The top 30 genes based on log fold change are labeled in the plot. (**C**) *Napsin A Aspartic Peptidase (Napsa) expression. Napsa* transcript was significantly higher in normal podocytes in the juxtamedullary (JM, pink) compared to the outer cortical (OC, green) zones. (**D**) Napsin A Aspartic Peptidase protein was validated by immunostaining showing a granular cytoplasmic pattern in proximal tubules of the JM. (**E**) *Vin Diagram*. Of 1,332 marker genes for the podocyte cluster 1249 (93.8%) were not differentially expressed between JM and OC (**F, G**) *Upset plots*. Upset plots show the number of genes in normal mice higher in the (JM) (**F**) and in the OC (**G**) unique to normal podocytes (POD), mesangial cells (MES) and glomerular endothelial cells (GEnC) and the number of genes that overlap between these glomerular cells. For normal podocytes, 585 and 470 genes were higher in the JM and OC respectively, of which 199 and 269 genes were unique to podocyte in the JM and OC respectively. (**H**) *Hallmark Pathways*. Hallmark pathway enrichment of DEGs higher in podocytes derived from the JM and OC ordered by the proportion enriched. The color of the dots indicates the statistical significance. k/K ratio represents the number of genes from the DEG genes that overlap with the total number of genes in that Hallmark pathway’s gene set.

To address whether podocyte defining genes differ between the regions, we examined expression of two lists of podocyte genes. First, we looked at the 1,332 marker genes for the podocyte cluster and found that the vast majority (93.8%) were not differentially expressed between JM and OC (**Figure 2E**). Secondly, by parsing the genes based on a list of 51 canonical podocyte genes we assembled (**Table 3**), only 3 were differentially expressed, *Magi1* and *Mapt* in the OC and *Nephronectin* (*Npnt*) in the JM. While many genes differed between JM and OC in podocytes, most genes that differentiate podocytes from other kidney cell types were consistent across both regions.

**Table 3.**
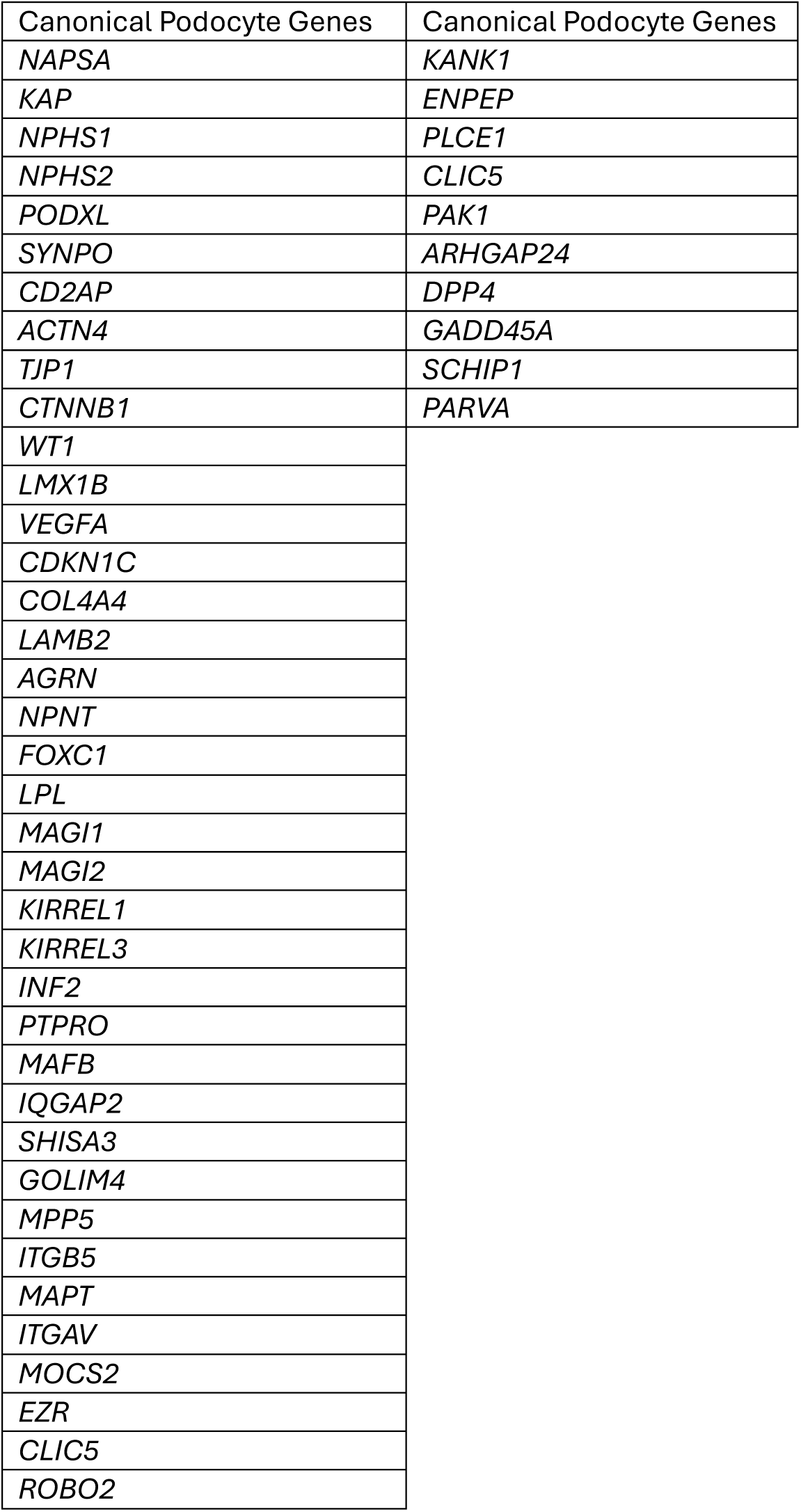

**Table 4.**

| *Lower at FSGS D7 |  | *Lower at FSGS D28 |  |
| --- | --- | --- | --- |
| *JM* | *OC* | *JM* | *OC* |
| Lpl<br>**Wt1** | Actn4<br>Enpep<br>Foxc1<br>Itgb5<br>Kank1<br>Lmx1b<br>Lp1<br>Nephs1<br>Nphs2<br>Shisa3<br>Vegfa<br>**Wt1** |  | **Wt1** |
| * *P* <0.05 vs. normal using *FDR* <0.05 |  |  |  |

We then questioned which of the differentially expressed genes were specific to podocytes and not present in two other cell types present in the glomerular tuft, glomerular endothelial cells (GEnC) and mesangial cells (MES). Following the same workflow as for podocytes (POD), we determined the significantly differentially expressed genes in the OC and JM for both cell types. Comparing those with those identified for the podocytes we determined that of the 470 genes that were expressed significantly higher in the OC, 269 were podocyte-specific (**Figure 2F**), similarly, of the 585 JM-enriched genes, 199 were podocyte-specific (**Figure 2G**).

To obtain insights whether these differential genes reflected differential pathways activated in OC *vs.* JM podocytes, we performed hypergeometric enrichment analysis using the Hallmark gene sets (**Figures 2H**). The most noteworthy observation of this analysis was that the top pathways enriched in JM podocytes were metabolic pathways such as oxidative phosphorylation, adipogenesis, fatty acid metabolism, and glycolysis. This was further corroborated when we specifically analyzed the genes in these pathways (**Supplemental Table S1**). Many of the genes in the Hallmark pathways for oxidative phosphorylation, glycolysis and fatty acid metabolism were expressed significantly higher in JM podocytes than in OC podocytes, while there were relatively few that were higher in normal JM than OC podocytes.

Together, these data support the notion that podocytes from the JM and OC region of the kidney share most defining gene characteristics, but also have some notable differences in their transcriptome.

### Region-Specific Podocyte DEGs in Response to Experimental FSGS

Next, we extended our analysis to include injured podocytes using the same pseudo-bulk approach. Volcano plots comparing the JM and OC regions at FSGS day 7 and at day 28 demonstrated distinct DEGs as shown in volcano plots in **Figures 3A-D**. Using upset plots to show trends in gene expression revealed a surprising heterogeneity in the injury response (**Figure 3E,F**).

**Figure 3:**
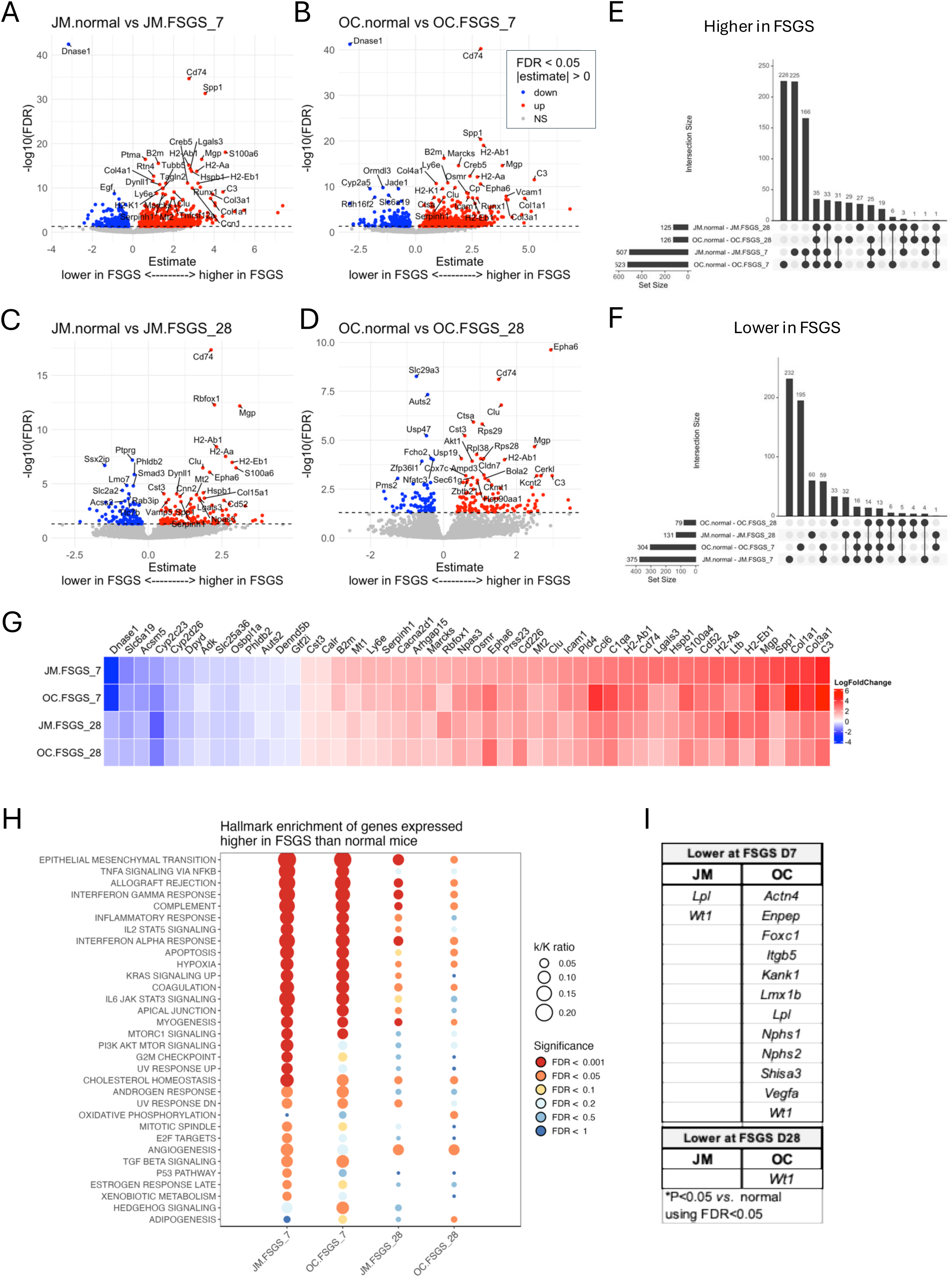
Podocyte’s Transcriptomic Response to Injury in Experimental FSGS. (A-D) *Volcano plots*. Volcano plots of the top 30 most enriched DEGs in podocytes lower (blue) and higher (red) in FSGS over normal at day 7 in the JM (**A**) and OC (**B**) and day 28 in the JM (**C**) and OC (**D**). All comparisons are from the labeled group compared to normal mice. Blue indicates genes expressed higher in normal mice; red indicates genes expressed higher in FSGS mice. **(E,F)** *Upset plots in FSGS by zone and time course*. The horizontal rows below each graph show the number of DEG’s in podocytes that are higher (**E**) and lower (**F**) in FSGS compared to normal.(**G**) *Heatmap by zone and FSGS time course* shows the limited number of DEGs shared throughout disease (day 7 and day 28) in the JM and OC. (**H**) *Dot plot.* Hallmark pathways in podocytes of the JM and OC at days 7 and 28 of FSGS ranked by significance (colors of dots shown on the right); the size of the dot (k/K ratio) represents the percentage of DEGs of the total number of genes in each given Hallmark pathway. (**I**) *Canonical podocyte genes*. Table showing podocyte genes significantly lower at each time point by zone. Wt1 (bold font) is the only overlapping podocyte gene that is lower in 3 of the 4 comparisons.

The most obvious trend was that the majority of DEGs (both up- and down-regulated) were between the normal mice and the mice at day 7 of FSGS, while the numbers of DEGs between normal and day 28 of FSGS were much smaller. This suggested that the peak of the transcriptomic changes in disease was at day 7 and that most of the recovery and/or repair was done by day 28. In fact, only a limited number of DEGs were shared throughout disease (day 7 and day 28) irrespective of location. These overlapping DEGs are shown in the heatmap in **Figure 3G**.

The other insight from this analysis was that there were many DEGs that were unique to the OC and the JM during FSGS day 7 and 28. Among the upregulated DEGs at day 7, 226 were OC-specific, 225 JM-specific and 166 shared between OC and JM. Similarly, in the downregulated DEGs 195 were OC-specific, 232 JM-specific, and only 59 shared between OC and JM. Interestingly, of the top 30 increased genes at FSGS day 7 several related to extracellular matrix changes with *Mmp2* and *Mmp12* in the OC and *Col1a1*, *Col3a1*, *Col5a3*, *Fn1* and *Mgp* in the JM. This region-specific response was also observed at day 28, albeit blunted due to the reduced number of DEGs present at this timepoint.

Next, to identify whether Hallmark pathways were differentially activated in OC *vs.* JM podocytes, we performed hypergeometric enrichment using the Hallmark gene sets (**Figures 3H**). This yielded several interesting findings. First, multiple pathways were enriched in the OC and JM podocytes at day 7. These primarily included inflammatory responses such as TNFa Signaling via NFkB, Interferon Gamma Response, IL2 Stat5 Signaling and Allograft Rejection. Second, most of these pathways were reduced by day 28 consistent with recovery and/or repair. This was seen for both regions, albeit more marked in the OC podocytes. We previously reported that podocyte depletion in both the OC and JM drives the development of injury in this model of FSGS.^14^ Indeed, this was confirmed in the enrichment, where Apoptosis, Epithelial Mesenchymal Transition and Apical Junction were upregulated early in the disease process at day 7 but tapered off at day 28.

Interestingly, when the individual genes of the Apoptosis gene set pathway were investigated, the response was not uniform but showed both shared and region-specific responses (**Supplemental Table S2**).

Finally, parsing the genes based on the list of canonical podocyte genes between JM and OC following injury at day 7 and day 28 (**Figure 3**). Most of these genes were down regulated in the OC at day 7, with only *Lpl* and *Wt1* impacted in the JM. Moreover, *Wt1* was the only gene that remained low in OC podocytes at day 28.

Taken together, the DEG results showed that following injury, there were differences in expression between podocytes in the JM and OC, and that injury was more severe at day 7 and significantly receded by day 28.

### Sub-clustering identifies the transition from healthy to injured podocytes

To investigate the injury response at the cellular level, we re-integrated and performed unsupervised sub-clustering of the 7,390 podocytes (**Figure 4A**). This identified 5 podocyte subclusters, which were characterized by unique gene expression patterns (**Figure 4B,C**). Using markers of canonical podocyte genes, parietal epithelial cell (PECs) and FSGS injury we assigned podocytes into healthy podocytes (cluster 0), PEC-like cells (cluster 1), and two populations characterized by reduced canonical gene expression, but absence of PEC and injury markers, that may represent cells transitioning from healthy to injury (clusters 2 & 3). Finally, cluster 4 represented injured cells (**Figure 4B**). Cluster 4 was characterized by low expression of canonical podocyte markers (*Podxl*, *Nphs1*, *Nphs2*, *Synpo*, *Cd2ap*) as well as genes known to be differentially expressed upon podocyte injury (*Havcr1*, *Serpine1*, *S100a6,C3*). It also expressed the JM markers (*Kap*, *Napsa*), suggesting that more of the injured podocytes in cluster 4 resided in the JM. This was confirmed by the OC/JM distribution plot, which demonstrated that while cluster 4 cells were derived from both regions, a higher percentage (17.3) were from the JM while only 6.9% were from the OC (**Figure 4D** and **Supplemental Figure S2C**). Furthermore, cluster 4 is primarily present in the day 7 podocytes, and its numbers drops to control levels at day 28 (**Figure 4E**). This is likely due the fact that by day 28, most of the extremely damaged podocytes have detached and/or undergone cell death and thus have been effectively eliminated them from the analysis. In line with the focal nature of our podocytopathy, it is not surprising that this cluster only represents a small proportion of the total podocytes (398 cells, 5.4%), yet at day 7 it comprises a significant number of 17.3% of the JM and 6.9% of the OC podocytes (**Figure 4D**). The top markers for each podocyte cluster are shown in **Figure 4C**. Finally, we parsed the individual clusters by sex, but did not detect any significant differences in cluster distribution by sex (**Figure 4G**) in line with our bulk mRNA analysis of aging podocytes.^34^

**Figure 4:**
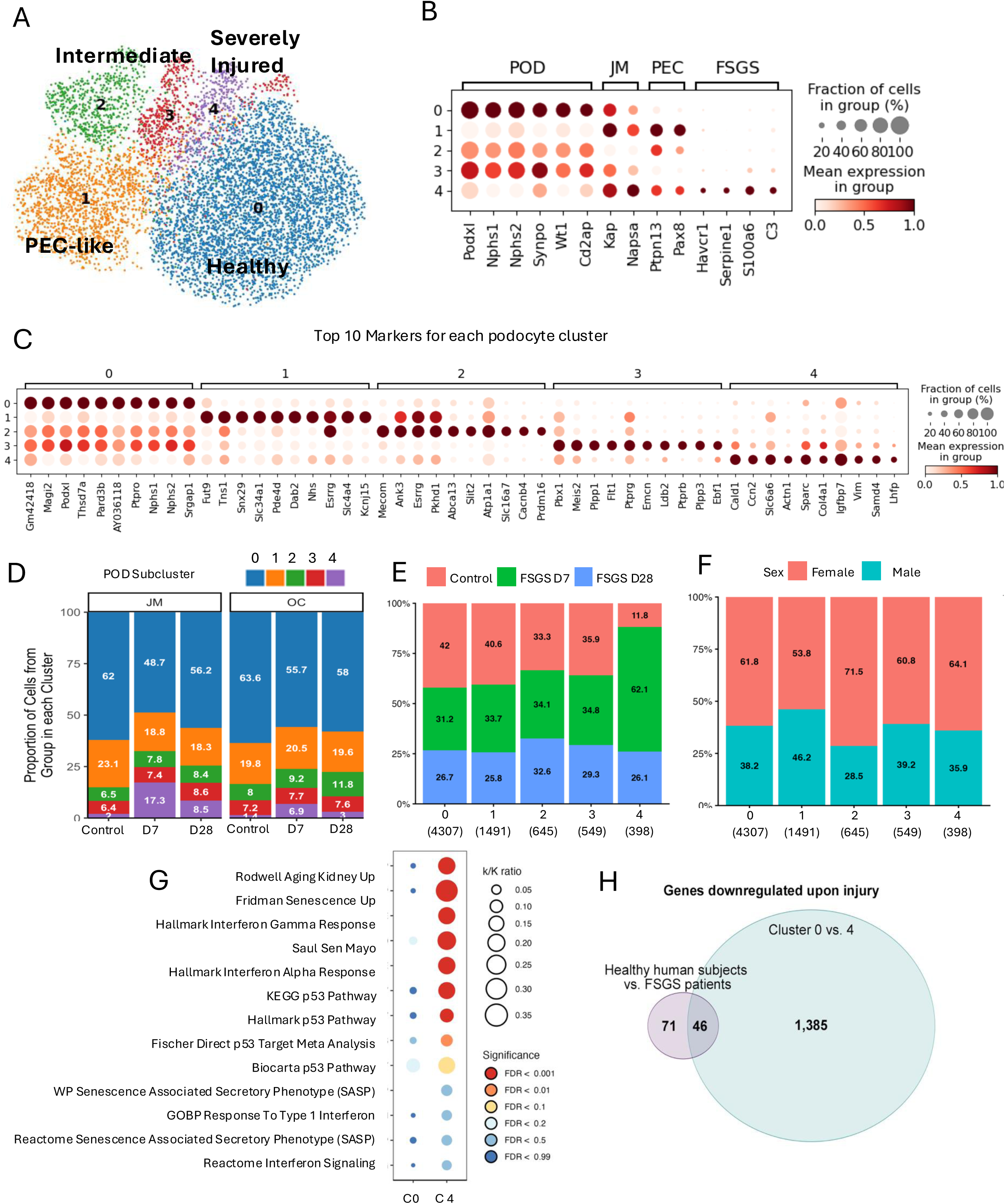
(A) *Sub clustering of Podocytes*. UMAP sub clustering of the 7,390 podocytes resulted in 5 distinct clusters – healthy (zone 0, blue), PEC-like (zone 1, orange), intermediate injury (zones 2 and 3, green) and severely injured (zone 4, purple). **(B)** *Dot plot* of expression of normal podocyte markers, JM markers, PEC markers, and a few of the top FSGS DEGs. The high expression of podocyte markers in cluster 0 suggests this is a cluster of normal healthy podocytes. The low expression of healthy POD markers and higher expression of FSGS DEGs in cluster 4 suggests that these are injured podocytes. **(C)** *Dot plot* of expression of the top DEG markers for each podocyte cluster. **(D)** *Stacked box plot* showing the percent of cells in each cluster arranged by disease time point and kidney zone (JM vs OC). The color represents each cluster and the white text in the bars is the percent of each cluster. **(E)** *Stacked box plot* showing the percent of cells in each cluster from each group at day 0 (pink), day 7 (green) and day 28 (blue) of FSGS. The color represents the group, the balck text in the bars is the percent and the number in parentheses under the cluster number represents the total number of cells in that cluster. **(F)** *Stacked box plot* showing the percent of cells in each cluster by sex Female (pink) and Male (blue). The black text in the bars indicates the percentage and the number in parentheses under the cluster number represents the total number of cells in that cluster. (**G**) *Hallmark Pathways.* Dot plot of select hallmark pathways in normal podocytes (subcluster 0, c0) compared to injured podocytes (cluster 4, c4). The color of the dots indicates the statistical significance. k/K ratio represents the number of genes from the DEG genes that overlap with the total number of genes in that Hallmark pathway’s gene set. (**H**) *Vinn Diagram.* Comparison of DEGs from the human patients with the DEGs of cluster 4 *vs.* cluster 0 mouse podocytes. Of the 71 downregulated genes in the podocytes of healthy patients compared with FSGS patients, 46 overlapped with the 1385 downregulated genes in cluster 0 (healthy) vs. cluster 4 (injured).

**Figure 5:**
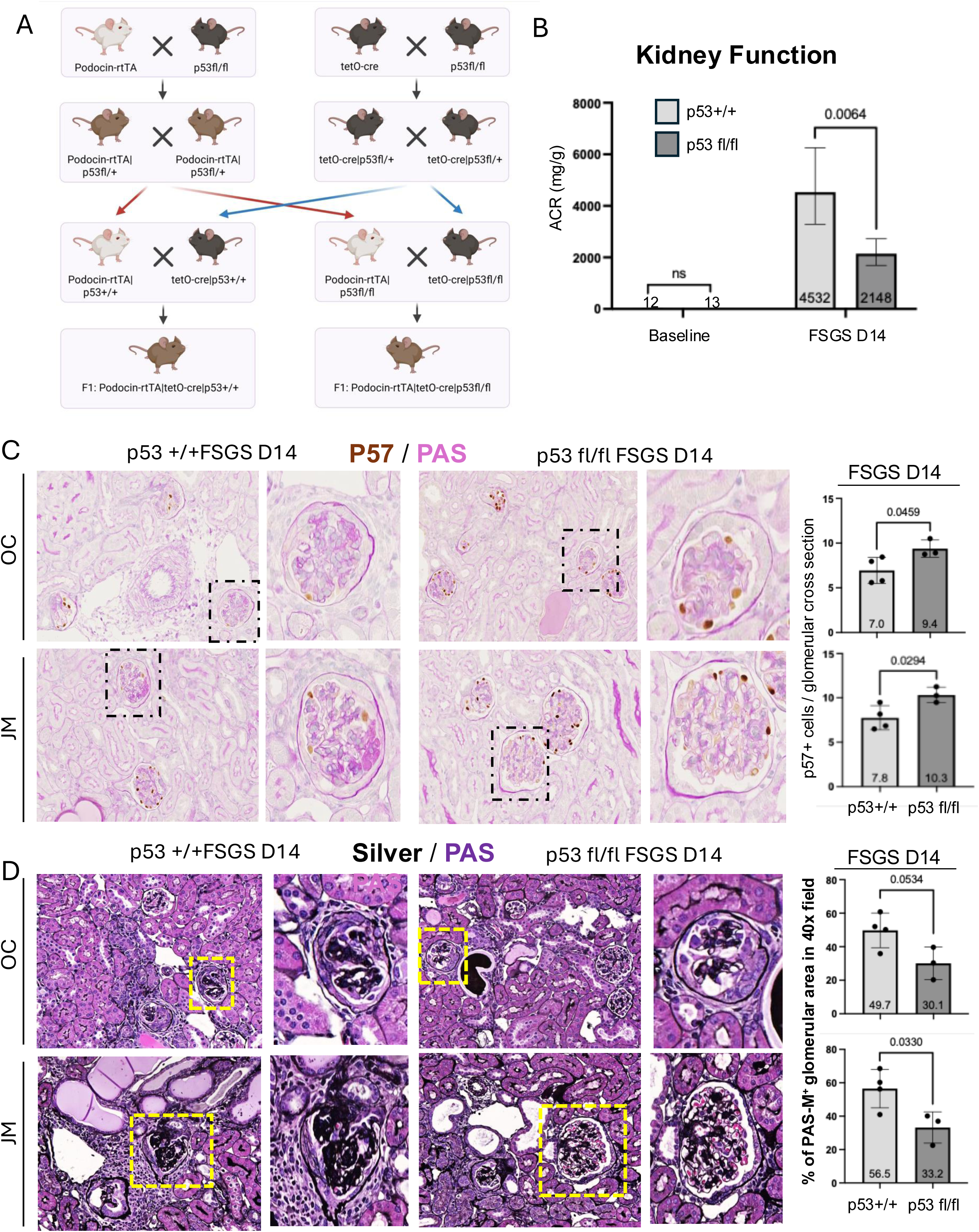
Podocyte and glomerular damage lower in podocyte-specific p53 mutant mice with FSGS. **(A)** *Mouse breeding* schema showing the crosses of Podocin rtTA, p53 fl/fl and tetO-cre mice used to establish mice with doxycycline inducible podocyte specific ablation of p53. (**B**) *Kidney function.* Bar Graph of albumin creatinine ratios(ACR) at baseline and day 14 of FSGS in p53 +/+ (light gray) and p53 fl/fl (dark gray) mice. ACR was higher in p53 +/+ compared with p53 fl/fl mice (**C**) *Podocyte number*. Representative images of p57 staining in the OC and JM and bar graphs of quantification of the number of p57 positive cells per glomerular cross section. Podocyte number was lower in p53 +/+ compared with p53 fl/fl mice (**D**) *Glomerulosclerosis.* Representative images of Jones silver staining in the OC and JM and bar graphs of quantification of the percentage of the glomerular area stained with silver. Silver staining was higher in p53 +/+ compared with p53 fl/fl mice indicating matrix expansion and more scarring.

To further compare gene expression of the injured podocytes, present in cluster 4 with their healthy cluster 0 counterparts, we compared the expression of the top DEGs (**Supplementary Figure S2A,B**). As expected, many of the canonical podocyte genes (e.g., *Nphs1*, *Nphs2*, *Podxl* or *Vegfa*) were present in the list of genes downregulated genes in the injured podocyte cluster 4 (**Supplementary Figure S2A**). While there was some variation between OC and JM in healthy podocyte cluster 0 with respect to the podocyte canonical genes, there was a more pronounced difference in their downregulation upon injury in the JM with the OC showing a less pronounced effect in cluster 4 (**Supplementary Figure S2A**). Similarly, injury-dependent genes, while conserved in their overall response, show some difference in the degree of the expression based on the region with some higher in the OC and others higher in the JM (**Supplementary Figure S2B**).

To get a better understanding of the transcripts upregulated in the injured podocyte cluster 4, we performed hypergeometric enrichment. We focused on cellular senescence/aging and inflammatory pathways that we recently discovered were altered in bulk mRNA-sequencing in response to injury to non-aged podocytes.^54^ We took a deeper dive into these pathways in the most injured podocyte subcluster (subcluster 4) compared with the healthy podocytes (subcluster 0). As shown in **Figure 4G**, senescence gene sets such as The Rodwell Kidney Aging Up, Fridman Senescence Up and Saul Sen Mayo were highly enriched; similarly, some inflammatory gene sets such as Hallmark Interferon Alpha Response or Hallmark Interferon Gamma Response were highly enriched. Surprisingly, gene sets for the senescence-associated secretory phenotype (i.e. WP Senescence Associated Secretory Phenotype SASP and Reactome Senescence Associated Secretory Phenotype SASP), were only modestly enriched.

Gene sets examining the known trigger for stress-induced senescence p53, such as Kegg p53 Signaling Pathway,^55^ Fischer Direct p53 meta-analysis pathway,^56^ and Hallmark p53 Pathway^57^ were all robustly increased in injured cluster 4 podocytes, and not healthy cluster 0 podocytes (**Figure 4G**).

Recently, Padvitski *et al.*^58^ reported on a podocyte damage score (PDS) to track damage during the progression of CKD. This score is driven by 42 genes, 39 genes that tend to be expressed lower in damaged podocytes (NEG) and 3 genes that tend to be expressed higher in damaged podocytes (POS). We examined the expression of these genes and found that the 39 NEG genes had the lowest expression in cluster 4 while the 3 POS genes had the highest expression in cluster 4, suggesting higher podocyte damage in cluster 4 (**Supplemental Figure S2C**). Using these 42 genes we calculated the PDS for the podocyte subclusters with cluster 0 yielding the lowest score and cluster 4 the highest (**Supplemental Figure S2D**). Interestingly, the other clusters (1-3) exhibited podocyte damage scores higher than cluster 0, but lower than cluster 4, supporting the conclusions that they are likely intermediary cell populations.

To investigate the translational potential of our data, we compared it to a recent study by Deleersnijder *et al.*,^59^ which investigated podocytes from human patients with primary FSGS and maladaptive FSGS by snRNA-seq. Comparing the DEGs from the human patients with the DEGs of cluster 4 *vs.*cluster 0 mouse podocytes showed that of the 71 genes downregulated in the podocytes of healthy patients compared with FSGS patients , 46 overlapped with the 1385 downregulated genes in cluster 0 (healthy) vs. cluster 4 (injured) (**Figure 4H**). The 46 overlapping genes are listed and compared in **Supplemental Figure 2E**.

### Ablation of p53 blunts the podocyte injury phenotype

Among the most interesting findings from the analysis of injured podocyte cluster 4 was the upregulation of p53 signaling. The expression of the transcripts for *Trp53* as well as the p53 transcriptional target *Serpine1*, were both increased in the podocyte cluster in the JM at day 7 of FSGS (**Supplemental Figure S3A,B).** To confirm this increase at the protein level, immunostaining of mouse kidneys that underwent the same podocyte injury model of FSGS, showed increased staining for p53 protein (**Supplemental Figure 3C**) and Serpine1 (**Supplemental Figure 3E**) in both OC and JM podocytes at day 14.

These results prompted us to test the hypothesis whether the increase in the p53-pathway in injured podocytes was biologically relevant. To this end, we generated an inducible podocyte-specific p53 null mice, where p53 was specifically deleted in podocytes upon the administration of doxycycline as described in detail in the methods section (**Figure 5A**). To induce podocyte injury, the same experimental model of FSGS was induced by the administration of 2 consecutive doses of cytopathic sheep anti-podocyte antibody as reported previously.^14, 29, 30^ At day14, urine was collected for kidney function measurements (albumin creatinine ratio, ACR) and kidneys were harvested for histological analysis. To verify that p53 was ablated in p53fl/fl kidneys, immunostaining for p53 and Serpine 1 protein was performed in p53fl/fl kidneys at day 14 of FSGS. Staining for both p53 and Serpine 1 were absent in both OC and JM podocytes following doxycycline administration, indicating p53 was ablated in podocytes, however both proteins were present in other kidney cells (**Supplemental Figure 3D,F)**.

As expected, ACR significantly increased at day 14 FSGS in p53 +/+ mice, however, the increase was significantly less in the p53 fl/fl mice where p53 was specifically deleted in podocytes (**Figure 5B**).

We have previously reported podocyte number, as measured by p57 staining, decreases following podocyte injury induced by the cytopathic anti podocyte antibody, with a mean starting number of 12.7 ± 0.9 podocytes per glomerular cross section.^54^ The results of the current study showed a similar decrease in podocyte number in p53 +/+ mice in both OC and JM at day 14.

While podocyte number also decreased in p53 fl/fl mice, the loss was significantly less in both the OC and JM (**Figure 5C**).

To assess changes in basement membranes and glomerular scarring, Silver Jones staining was performed. Indeed, an increase in the % of the glomerular area staining for silver was significantly increased in p53 +/+ mice in the OC and even more so in the JM at day 14. Again, the % of the glomerular area staining also increased in p53 fl/fl mice, however the increase was significantly less in both the OC and JM (**Figure 5D**)

We have recently reported on the induction of the senescent podocyte phenotype in our experimental model of FSGS.^54^ In order to determine if p53 plays a role in this phenotype, immunostaining for the senescent maker SA-b-galactosidase (SA-b-gal), p16 as a marker of the p16/Rb axis of senescence and p21 as a marker of the p53/p21 stress-induced axis of senescence was performed. The results of the current study showed similar increases in staining for SA-b-gal in podocytes in p53 +/+ mice in both OC and JM (**Figure 6A**). SA-b-gal staining did increase in PECs in p53 fl/fl mice, but podocyte staining was much less compared to p53 +/+ mice (**Figure 6B**). Staining for p16 was pronounced across many cell types, including podocytes in p53 +/+ mice in both the OC and JM regions (**Figure 6C**), however, while staining was also pronounced in p53 fl/fl mice, it was noticeably less in glomeruli and particularly podocytes (**Figure 6D**). Finally, staining for p21 was notable in podocytes in p53 +/+ mice in the OC and even more so in the JM (**Figure 6E**), but staining was largely absent in p53 fl/fl mice (**Figure 6F**).

**Figure 6:**
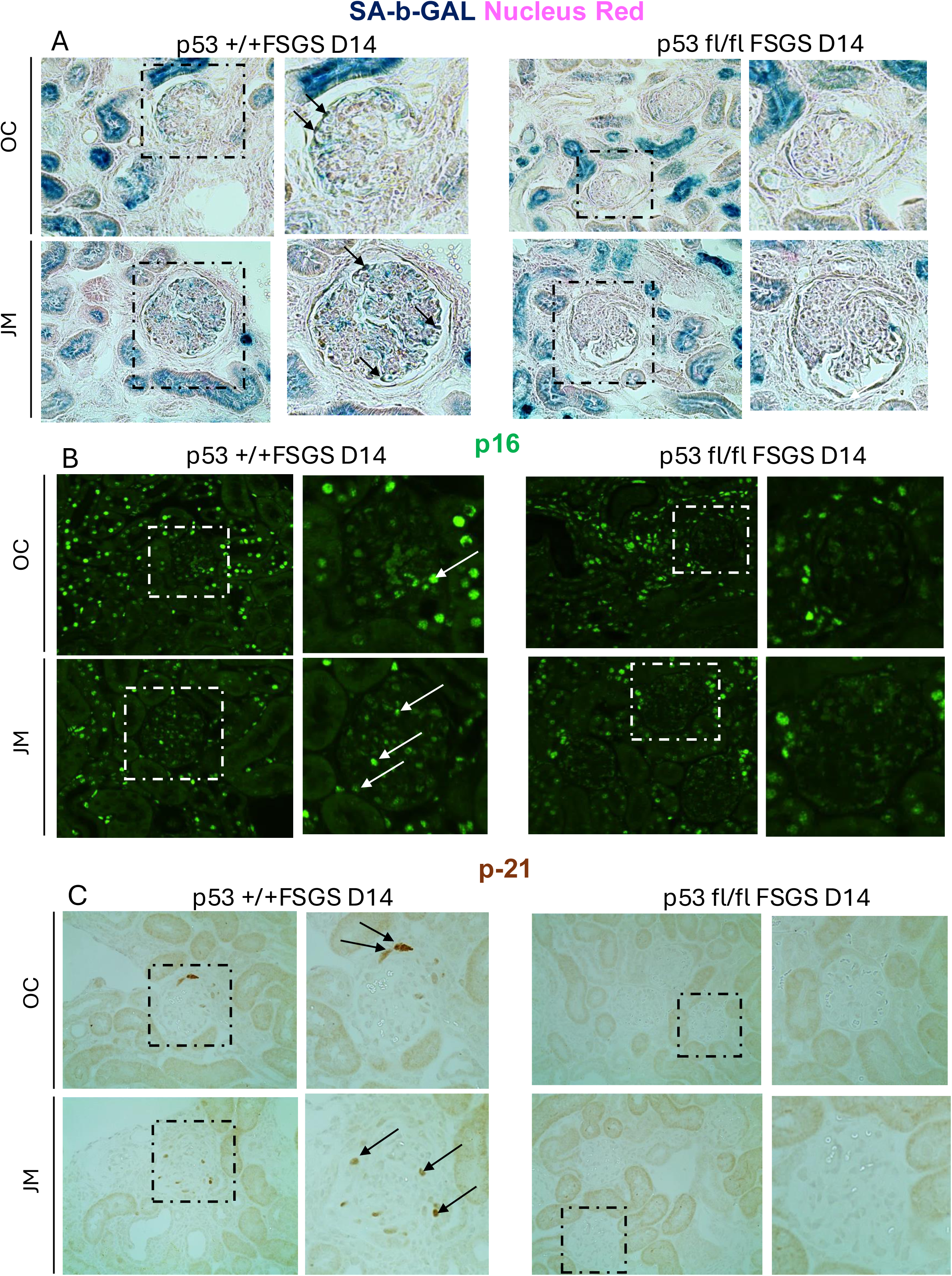
Senescent podocyte phenotype lower in podocyte-specific p53 mutant mice with FSGS. (**A)** *SA-ß-Gal staining*. Representative images of SA-ß-Gal staining in the OC and JM of p53 +/+ (left) and p53 fl/fl (right) FSGS mice shows decreased senescence following p53 ablation (arrows indicate examples) (**B**) *p16 staining*. Representative images of p16 staining in the OC and JM of p53 +/+ (left) and p53 fl/fl (right) FSGS mice shows decreased p16 staining in podocytes of p53 fl/fl compared to p53 +/+ mice (arrows indicate examples), but not in other cells following p53 ablation (**C**) *p21 staining*. Representative images of p21 staining in the OC and JM of p53 +/+ (left) and p53 fl/fl (right) FSGS mice shows decreased p21 staining in podocytes following p53 ablation compared with p53 +/+ mice (arrows indicate examples).

In summary, these results showed that the increase in p53 in the injured podocyte population contributes significantly to a decrease in kidney function, loss of podocytes, increased scarring and a senescent podocyte phenotype and these changes were significantly reduced when p53 expression was specifically ablated in podocytes.

### Changes to Adhesion Genes and Urinary Podocyte Transcriptional Changes

As the injured podocytes from cluster 4 were primarily present at day 7 and returned to pre-injury levels by 28 days (**Figure 4D**), we wondered if podocyte detachment was responsible for this shift. We therefore looked for changes in adhesion/de-adhesion genes, comparing injured to healthy podocytes (i.e. cluster 4, injured vs cluster 0, healthy) using a panel of genes reported to be important in podocyte adhesion under healthy and injured conditions.^60, 61^ As shown in the dot plot in **Figure 7A**, no clear trends were discernable. On one hand, cluster 4 exhibited a decrease in podocyte adhesion genes such as collagens (*Col4a3*, *Col4a4*), canonical podocyte genes (*Cd2ap*, *Nphs1*, *Nphs2, Podxl* and *Slit2*), *Epb41l5*, which is involved in podocyte extracellular matrix assembly,^62^ as well as *Srgap1*, which is involved in foot process maintenance.^63^ On the other hand, other genes favoring podocyte adhesion such as the tetraspanin *Cd151*,^64^ dystroglycan *Dag1*,^65^ the diphthamide biosynthesis genes *Dph1*, *Dph2* & *Dph3*,^61^ integrin (*Itga2*, *Itga3*, *Itgav*, *Itgb1* and *Itgb3*) and Laminin 521 (*Lama5*, *Lamb2* and *Lamc1*) were increased.

**Figure 7:**
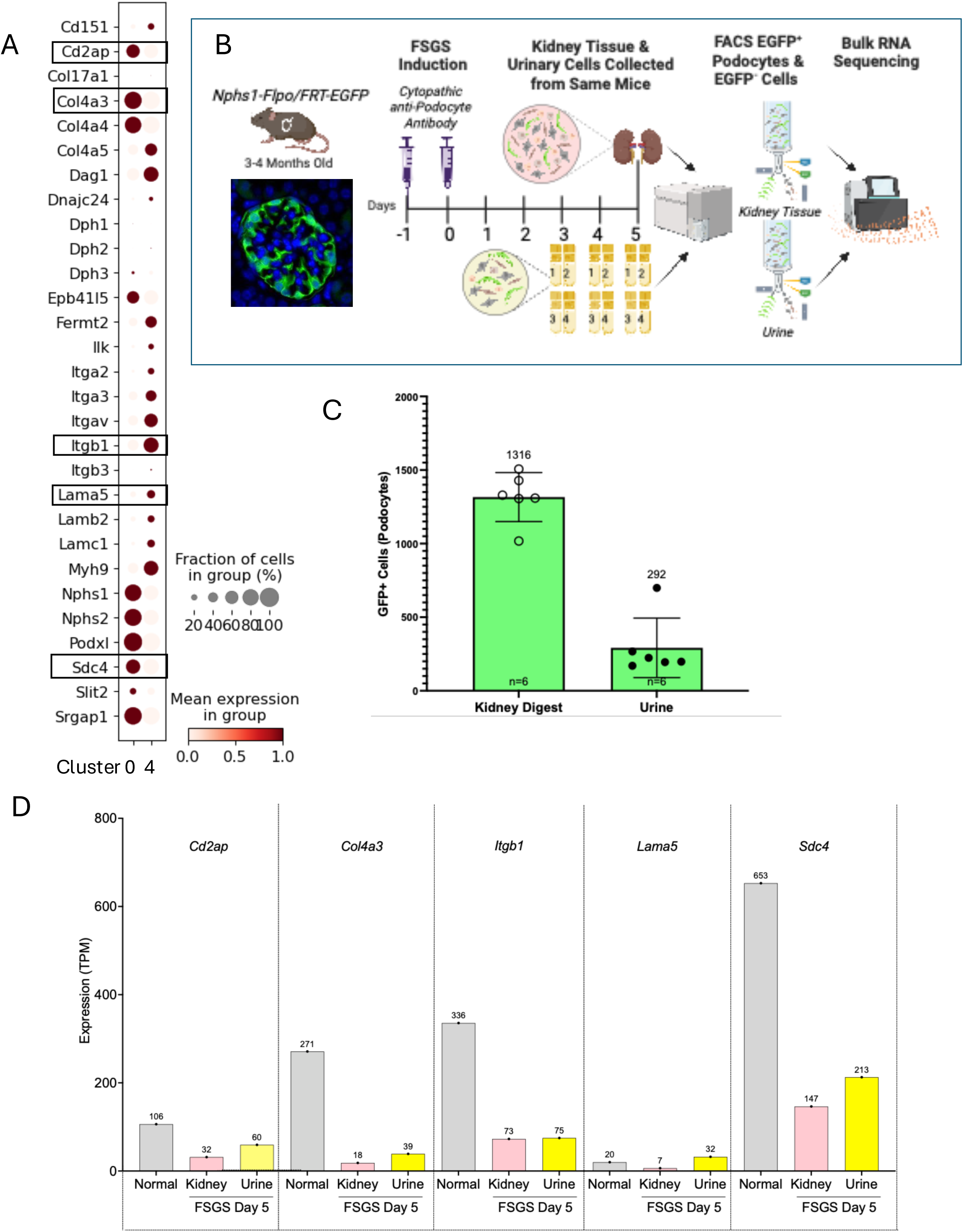
Changes to Adhesion Genes and Urinary Podocyte Transcriptional Changes. (**A**) *Podocyte adhesion genes in healthy and injured conditions.* Dotplot of select podocyte adhesion genes under healthy and injured conditions in normal podocytes (subcluster 0, c0) compared to injured podocytes (cluster 4, c4). The color of the dots indicates the mean expression. The size of the dots indicates the percentage of cells expressing in each cluster (0 vs. 4) (**B**) *Experimental Design*. Glomerular cross section demonstrates podo-specific expression of the EGFP reporter in 4-month-old *Nphs1-FLPo/FRT-EGFP* reporter mice. Mice were injected with two injections of sheep cytopathic anti-podocyte antibody. Urines from each mouse were collected. In the same individual mice kidneys will be removed on days 5. Paired urinary and kidney cell pellets underwent FACS analysis to isolate EGFP^+^ podocyte and EGFP^-^ kidney cells and bulk mRNA sequencing was performed. (**C**) Graph of mean number of EGFP^+^ podocytes from urine and kidney from 6 different mice post FACS sorting. (**D**) Bar graph of the 5 adherence genes (*Cd2ap*, *Col4a3*, *Itgb1*, *Lama5* and *Sdc4)* that appeared in the bulk mRNA sequencing of matched kidney (pink) and urinary podocytes (yellow) that dethatched early at day 5 of FSGS. Gray bars show normal gene expression. Four genes followed the same trend as cluster 4 injured podocytes.

Thus, to more directly address podocyte detachment into the urine, we collected urine from *Nphs1-FLPo/FRT-EGFP* reporter mice where all podocytes are constitutively labeled with enhanced green fluorescent protein (**Figure 7B**). We treated with the cytopathic anti-podocyte antibody and collected urine on days 3-5, four times daily every 2 hours (**Figure 7B**). Pooling multiple spot collections yielded an average of 292 EGFP^+^ podocytes per mouse from the urine (**Figure 7C**). On day 5, kidneys were digested into single cell suspensions from the same corresponding mice and urinary and kidney digest cell pellets underwent FACS to isolate podocytes which yielded an average of 1316 EGFP^+^ podocytes (**Figure 7C**). EGFP^+^ and EGFP^-^(negative control) cells from the urine and kidney digest underwent bulk mRNA sequencing as we have reported (**Figure 7B**).^34, 48, 52, 66^

Of the adherence genes listed in **Figure 7A**, 5 appeared in the bulk mRNA sequencing of matched kidney and urinary podocytes at day 5 of FSGS. Besides *Itgb1*, which increased in cluster 4, the other 4 genes (*Cd2ap*, *Col4a3*, *Lama5* and *Sdc4*) followed the same trend as cluster 4 injured podocytes (**Figure 7D**).

## DISCUSSION

The different impact of the glomeruli located in the OC and JM to health and disease have been considered to reflect differences in blood flow, oxygen content, the size of the nephron and the species. By performing transcriptomic analysis on single nuclei from dissected OC and JM regions of normal healthy mice and mice with experimental podocyte injury, we augment this knowledge by addressing region-specific differences in podocyte gene expression under healthy and disease conditions.

The most compelling finding of the study were the transcriptional differences between podocytes of the JM and the OC. Using an FDR of < 0.05 in normal mice, we identified 1,055 DEGs between podocytes from the OC and JM, with 470 and 585 genes upregulated respectively. In practical terms, the identification of *Napsa*, a gene unique to the JM, provides a tool to molecularly distinguish the OC from JM. Yet mechanistic insights into the differences of OC and JM podocytes do not seem to come from the core podocyte program. Only 3 of 51 canonical podocyte genes were differentially expressed by region; *Magi1* and *Mapt* had higher expression in the OC and *Npnt* was expressed higher in the JM. In fact, the differences appear to be primarily metabolic in nature. A hallmark pathway analysis showed that in healthy mice JM podocytes were enriched for oxidative phosphorylation, glycolysis, and fatty acid metabolism compared to OC podocytes. Of the 50 most significant DEGs in the JM and in the OC, 8 and 10 respectively were in the oxidative phosphorylation pathway. While we have not yet functionally addressed this, a study by Brinkkoetter elegantly showed that anaerobic glycolysis and not oxidative phosphorylation is the major energy source of healthy podocytes.^67^ Thus, it will be interesting to re-examine their findings in respect to glomerular localization.

A similar scenario applied to injured podocytes, which showed many differential gene expression in the OC *vs.* the JM. Using an experimental mouse model of podocyte-depletion induced FSGS we have extensively reported on^14, 29, 30^ and similar to the Adriamycin model of FSGS,^22–24^ this model results in an earlier and more severe podocyte depletion in JM glomeruli compared to OC glomeruli.^14^ Among the DEGs, as expected, were the downregulation of canonical podocyte genes. This decrease was primarily seen in OC podocytes and had largely recovered by 28. The podocyte master transcriptional regulator *Wt1* was the only podocyte gene that was lower at both time points and in both OC and JM. This downregulation of canonical podocyte function was accompanied by an enrichment in inflammatory pathways (e.g. TNFa, interferon gamma and alpha, IL2/stat5, Kras, IL6/JAK/Stat3) in the podocytes of both the JM and OC early in disease at day 7. These were mostly lower by day 28 but were higher in JM podocytes compared to OC podocytes.

Among one of the features of the injury response in this model and a major cause of podocyte depletion is - as we have reported - apoptosis.^68–72^ Thus, it is not surprising that the apoptosis gene set was enriched early in disease coinciding with the abrupt loss of podocytes. The interesting part of this, however, is that although several apoptotic genes overlapped between JM and OC at day 7 (*Clu*, *Dap*, *F2R*, *Hspb1*, *Ifitm3*, *Lgals3*, *Mmp2*, *Tgfb2*, and *Top2a*), several other apoptosis-related genes were uniquely enriched in the JM or OC podocytes. Only *Clu*, *Hspb1* and *Lgals3* overlapped at all time points and zones. Clusterin (Clu) and heat shock protein 27 (Hspb1) promotes survival in podocytes in diabetic nephropathy.^73, 74^ Intracellular Galectin-3 (Lgal3) promotes cell survival whereas extracellular Lgal3 promotes apoptosis in other cell types under stress conditions.^75^ In the future, it will be interesting to see whether these changes constitute different apoptotic triggers, different aspects within the cell death program, or a different cellular response to the induction of apoptosis.

Our single nuclear RNA-sequencing analysis revealed 5 podocyte subclusters. Among those, cluster 0 represented normal, healthy podocytes with a high expression of canonical podocyte genes *Podxl*, *Nphs1*, *Nphs2*, *Synpo*, *Wt1* and *Cd2ap*. Cluster 1 was PEC-like, as it exhibited a lower expression of canonical podocyte genes, but expressed genes present in PECs such as *Ptpn13* and *Pax8*. Interestingly, PECs have the potential to trans-differentiate into podocytes and replace podocytes lost by injury or aging.^76–78^ Yet, whether cluster 1 cells are intermediates of this process still will need to be determined. Cluster 4 contained the smallest number of cells and was classified as the “podocyte injury” subcluster, typified by low/absent expression of canonical podocyte genes and the enrichment of transcripts that we have identified in our pseudo-bulk data as upregulated in FSGS (e.g., *Havcr1*, *Serpine1*, *S10016*, *C3*). This was substantiated in two ways: (1) When we compared our data to a recent report on a podocyte damage score,^79^ subcluster 4 reflects the most severely injured podocytes. (2) Comparing recent snRNA-seq data of human forms of FSGS with our data demonstrates significant overlap between the two data sets. Moreover, cluster 4 follows the disease trajectory identified by the pseudo-bulk analysis that injury was most prominent at day 7 and partially resolves by day 28.

Finally, the last two podocyte sub-clusters, clusters 2 and 3, appear to be intermediate states between normal podocyte cluster 0 and the injured podocyte cluster 4. Like in the case of cluster 1 future experiments using e.g., lineage tracing are needed to firmly establish this relationship.

A deeper dive into the injured podocyte cluster 4 showed that injured podocytes were enriched in gene sets associated with senescence and SASP, consistent with our previous report that a subset of injured podocytes acquired an aged/SASP phenotype in experimental mice and under cell culture conditions.^54^Yet, the more novel discovery was the robust activation of p53 stress-induced senescence pathways. We have previously shown that p53 signaling is activated upon podocyte injury,^47, 66^ and there was robust activation of three p53 pathways, many known transcriptional targets (e.g., *Cdkn1a*, *Mdm2*, *Trp5*, *Ccnd2*, *Ccnd3*, or *Serpine1*) and immunostaining for p53 in this study, along with its downstream targets p21 and Serpine1, suggesting that it plays a key role in the podocytes response to injury in FSGS.

While we previously demonstrated that inflammatory pathways such as PD1 or Nlrp3 are involved in podocyte aging and injury,^48, 52^ no data are yet available on the effect of master regulators of senescence. To this end, the ablation of p53 and the blunting of stress-induced senescence, is a first of its kind. In fact, using the same podocyte injury model of FSGS that we used for the snRNA-seq study, our results demonstrate that compared to injured wildtype mice, deletion of p53 specifically from podocytes was beneficial; besides inhibiting stress-induced senescence and bona fide targets of p53 (i.e., *p21* and *Serpine1*) it lowered albuminuria, improved podocyte number and lowered glomerulosclerosis,. Taken together these results show that the increase in p53 in the injured podocyte population was a major contributor to glomerular damage.

Podocyte depletion follows their detachment into the urine in clinical FSGS detachment into the urine.^55–58^ We did not examine their viability as we have previously reported in other models.^59, 60^ But, our data also support podocytes detachment contributing to podocyte depletion. The injured cluster 4 podocytes exhibited altered expression patterns when genes involved in podocyte adhesion to the glomerular basement membrane were looked at. Thus, to more thoroughly investigate this is warranted.

We also recognize some limitations of our study. (i) As with any single cell transcriptomic study, there are intrinsic limitations. Relating mRNA expression to protein expression is not linear. Complicating this even further, performing snRNA-seq over single cell transcriptomics only analyzes the pre-mRNA, which is basically the pool of actively transcribed genes, but does not address stable transcripts such as most housekeeping genes.^80, 81^ Thus, validation of key findings at the protein level is critical. (ii) We acknowledge that there are differences OC and JM glomeruli exist between humans and mice. Thus, our data in mice may not completely reflect the situation present in humans. (iii) We have only examined one model of experimental podocytopathy, and generalization of the results will require different model of podocyte injury.

In sum, our study identified major transcriptional differences in cortical *vs.* juxtamedullary glomeruli. This will greatly impact studying kidney function and its disruption during disease. The study highlights the need to perform the analysis of podocyte (and likely other cell types) by examining the OC and JM compartments independently whenever possible. This will avoid that avoid that the smaller number of JM podocytes is fractionally diluted by the larger number of OC podocytes. Finally, in the future generating podocyte-specific mutants or transgenic mice selectively targeting OC and JM podocytes will be critical to directly address the impact of these two podocyte populations. Finally, recognizing the differences between OC and JM podocyte will have clinical implications as therapeutic approaches may have to differ depending on which compartment is most disease relevant.

## Conflicts of Interest

The authors declare no potential conflict of interests.

## Grant Support

S.J.S. and O.W. were supported by National Institutes of Health (NIH) Grants 5R01DK056799-10, 5R01DK056799-12, 1R01DK097598-01A1, and UC2DK126006-2 as well as Department of Defense Grant PR180585/PR180585P1.

---

**Supplemental Figure S1:**
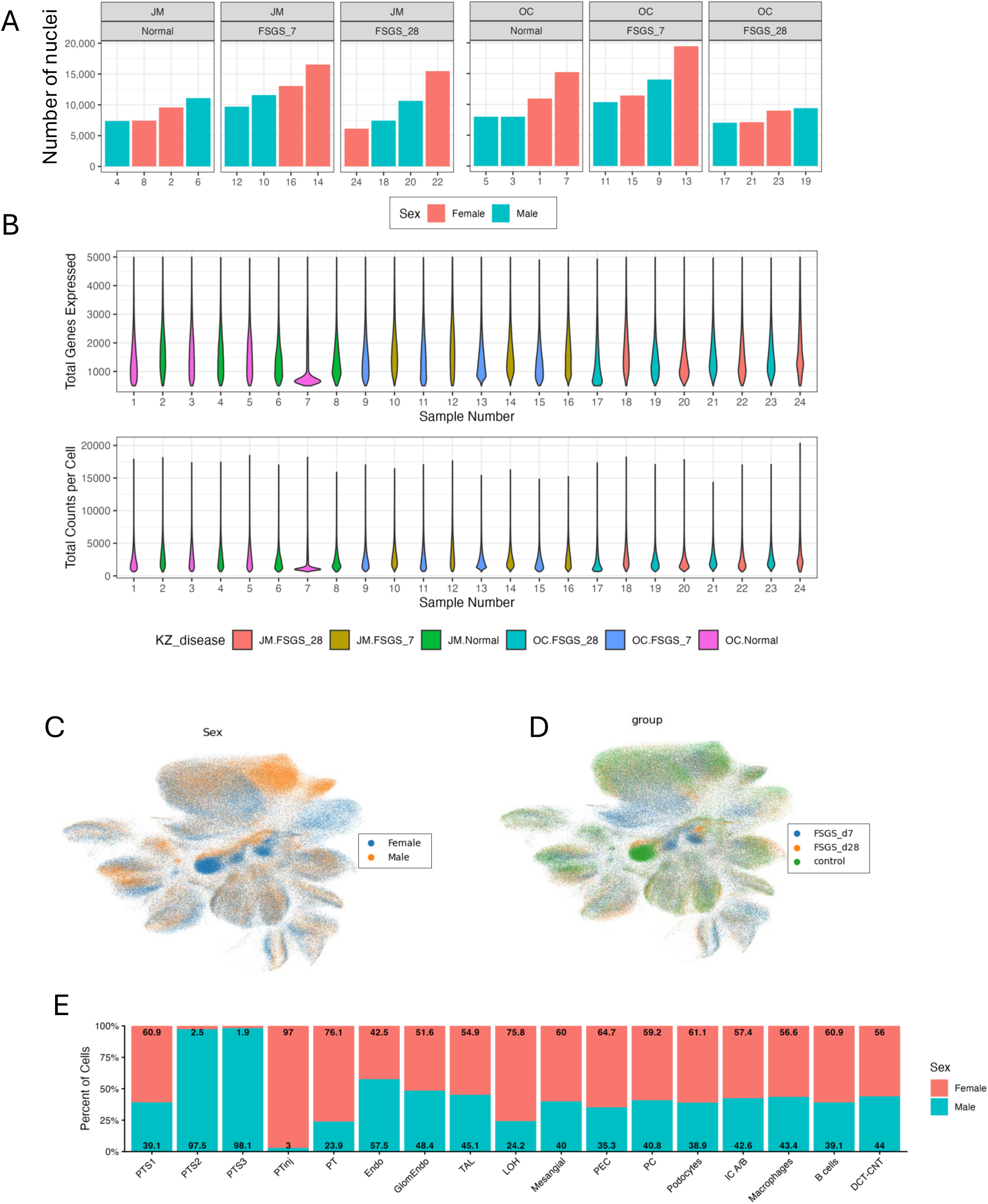
QC and Cell Distributions. **(A)** *Number of nuclei per sample.* Bar graphs show the number of nuclei obtained for each sample after filtering and processing by region (top), FSGS time point (top) and sex, Female (pink) and Male (Green). A total of 256,126 nuclei were obtained. **(B)** *Total genes expressed and counts per nuclei.* Violin graphs show the number of genes expressed (top panel) and the total counts per nuclei (bottom panel) for each sample after filtering and processing by region and FSGS time point. Sample 7 was omitted. (**C**) *Cluster distribution by sex.* UMAP shows the distribution of the 18 clusters by sex Female (blue) and Male (orange). Except for the proximal tubule populations (top right), the other cell types were relatively evenly distributed by sex across the clusters. (**D**) *Cluster distribution by disease progression.* UMAP shows the distribution of the 18 clusters between t=0 (control, green), FSGS day7 (blue) and FSGS day 28d (orange). Except for the proximal tubule populations (top right), the other cell types were relatively evenly distributed. (**E**) *Cluster percentages by sex.* Bar graph of the percentage of the cells in the 18 clusters by sex Female (pink) and Male (blue). Except for PTS2 (column 2), PTS3 (column 3), PT3_Inj_ (column 4), PT (column 5), and LOH (column 9) populations, the other cell types had similar percentages of cells in each cluster when filters for sex.

**Supplemental Figure S2:**
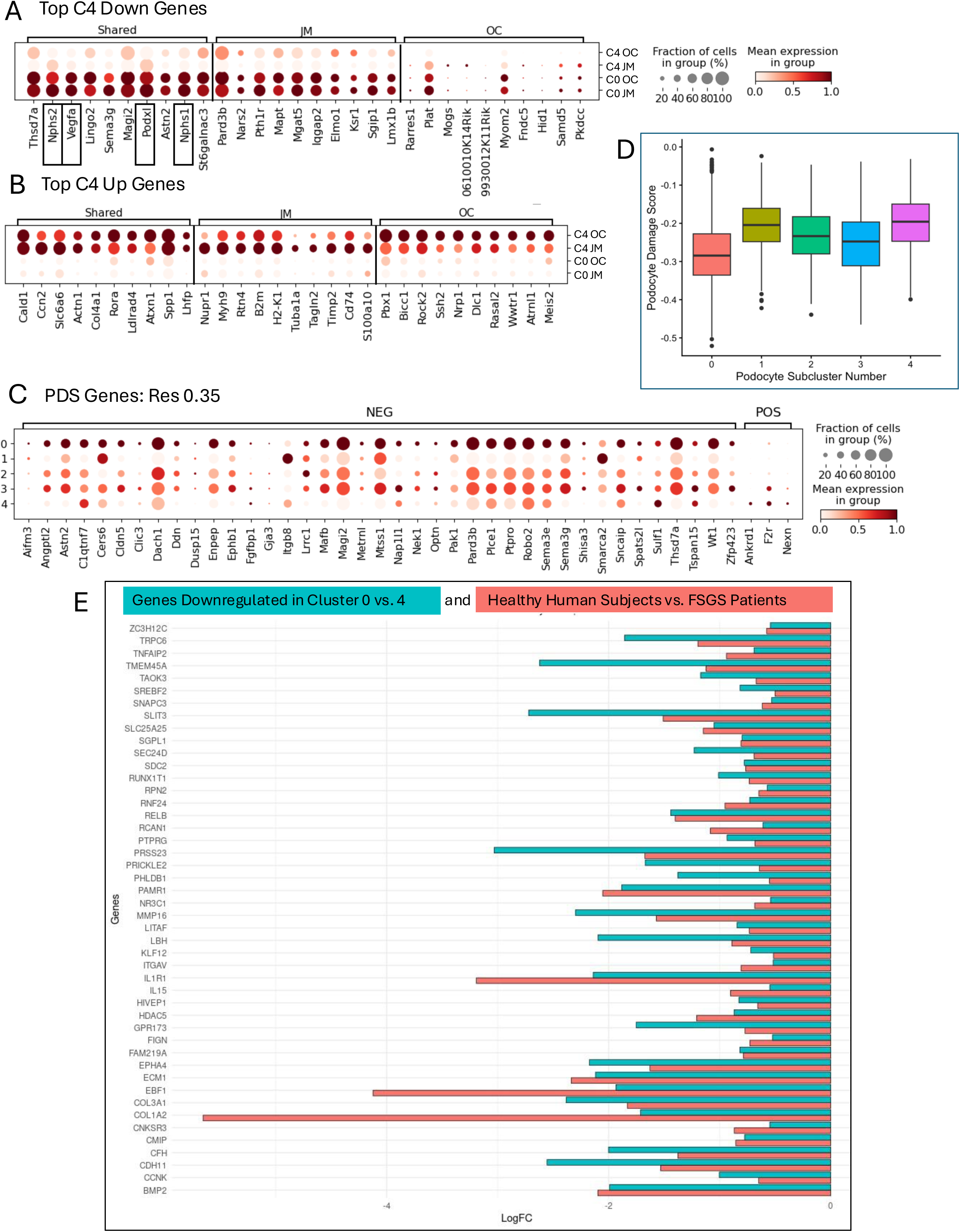
Additional characterization of podocyte subclusters. *Top differentially expressed genes in podocyte cluster 4*. **(A)** Dotplot showing expression of the top DEGs increased in cluster 4 injured podocytes compared to cluster 0 normal healthy podocytes organized and indicated by JM (middle), OC (bottom) or shared (top) regions. **(B)** Dotplot showing expression of the top DEGs decreased in cluster 4 injured podocytes compared to cluster 0 normal healthy podocytes organized and indicated by JM (middle), OC (bottom) or shared (top) regions. The color of the dots indicates the mean expression, and the size of the dots represents the percentage of podocytes in each cluster. Canonical podocyte genes (*Nphs1*, *Nphs2*, *Podxl* or *Vegfa*) were downregulated in the injured podocyte cluster 4 in the OC and were more decreased in JM regions. (**C**) *Podocyte damage score (PDS).* Dot plot showing expression of 42 genes that make up the PDS in each the 5 podocyte subclusters (0-4), 39 genes lower in damaged podocytes (NEG) and 3 genes higher in damaged podocytes (POS). The color of the dots indicates the mean expression, and the size of the dots represents the percentage of podocytes in each cluster. The 39 NEG genes had the lowest expression in cluster 4 while the 3 POS genes had the highest expression in cluster 4. (**D**) *PDS for podocyte subclusters.* Bar graph of PDS generated for podocyte subclusters 0 (red), 1 (moss), 2 (green), 3 (blue) and 4 (pink). Cluster 0 had the lowest score and cluster 4 the highest. Clusters (1-3) had podocyte damage scores higher than cluster 0 but lower than cluster 4. (**E**) *Mouse to human translational potential of our data.* Bar graph comparing and listing the 46 DEGs from downregulated in the podocytes of healthy patients compared with FSGS patients that overlapped with the DEGs that were lower in cluster 0 (healthy) vs. cluster 4 (injured) mouse podocytes.

**Supplemental Figure S3:**
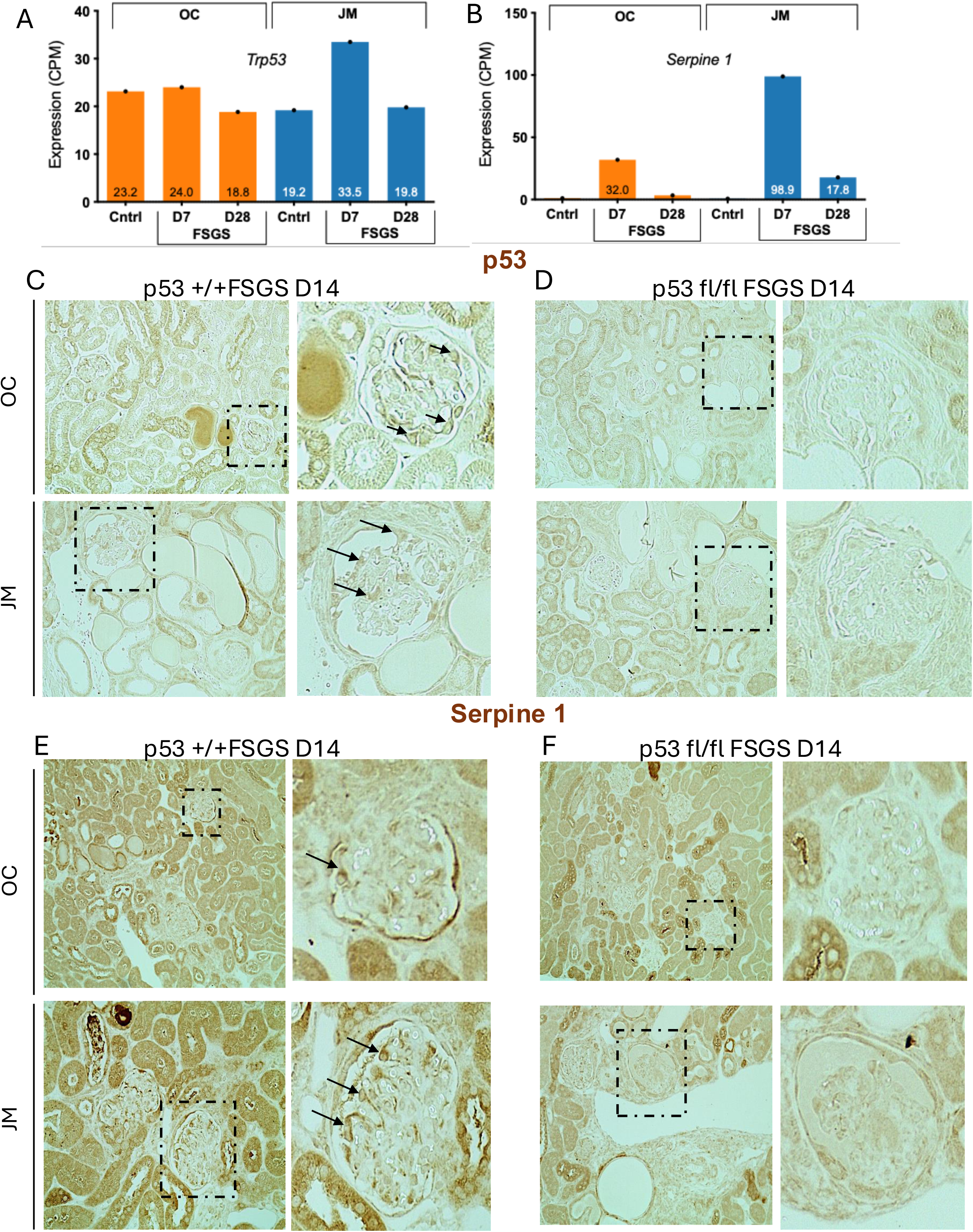
p53 and Serpine1 expression (A &. **B)** *Trp53 and Serpine1 transcript expression.* Bar graphs showing pseudo bulk analysis for the expression of *Trp53* and *Serpine1* in the podocyte cluster by OC (orange) and JM (blue) regions and by disease progression. Both transcripts were increased in the podocyte cluster in the JM at day 7 of FSGS (**C)** *p53 and Serpine1 protein*. Representative images of immunostaining for p53 protein in p53+/+ mouse kidneys at day 14 of FSGS. Staining for p53 increased in both OC and JM podocytes (arrows). **D)** Representative images of immunostaining for p53 protein in p53fl/fl kidneys at day 14 of FSGS. Staining for p53 was absent in both OC and JM podocytes, when p53 was ablated in podocytes but was present in other kidney cells. (**E**) Representative images of immunostaining for Serpine 1 protein in p53+/+ mouse kidneys at day 14 of FSGS. Staining for Serpine 1 increased in both OC and JM podocytes (arrows) and was particularly notable in PECs. (**F**) Representative images of immunostaining for Serpine 1 protein in p53fl/fl kidneys at day 14 of FSGS. Staining for Serpine 1 did not increase in OC and JM podocytes when p53 was ablated in podocytes but was present in other kidney cells.

**Supplemental Table S1:**
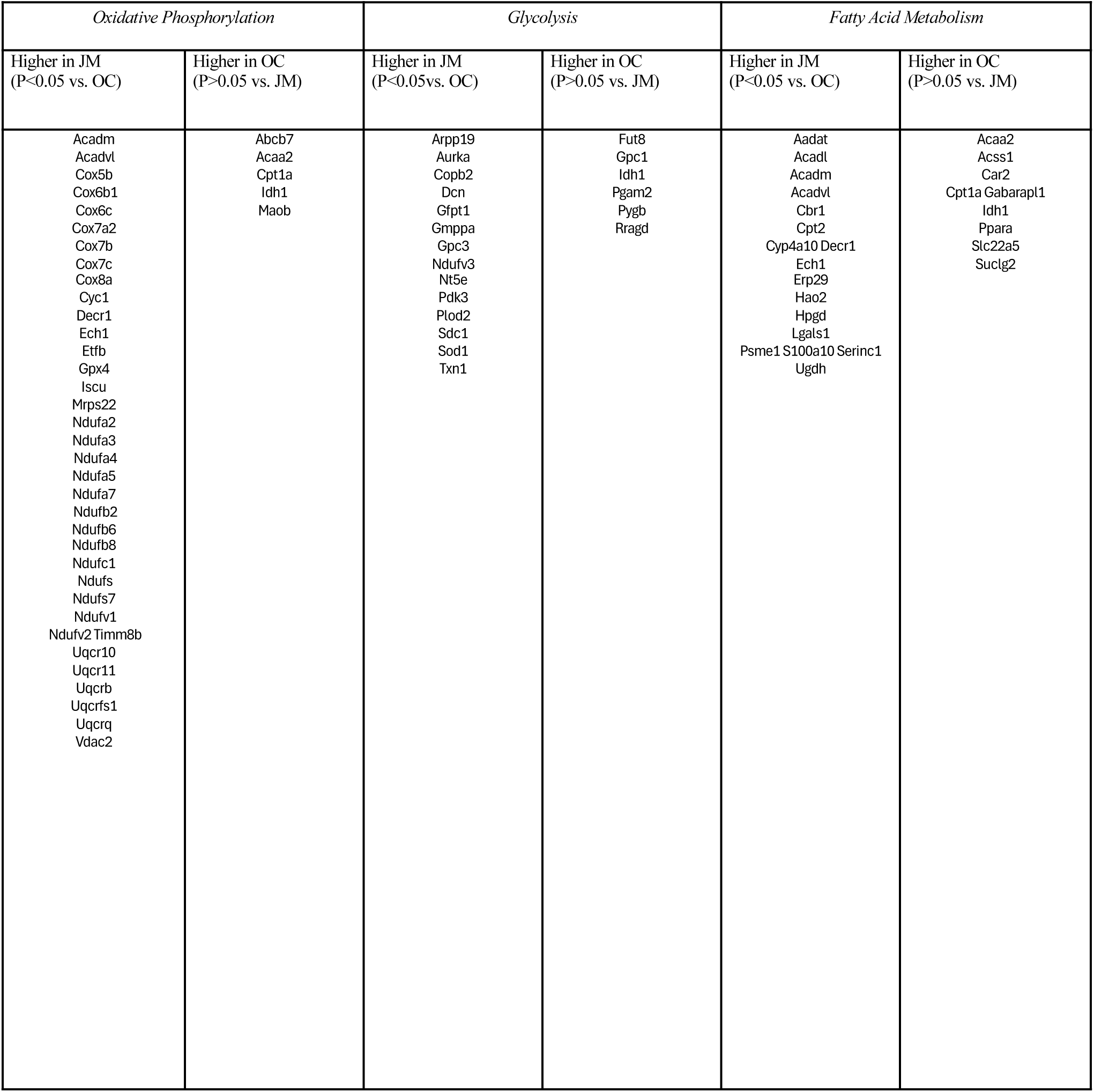
DEGs in major energy producing pathways comparing the OC *vs.* JM of normal, healthy podocytes.

**Supplemental Table S2.** Day 7 FSGS – Enriched Apoptosis Genes.

| **Day 7 FSGS – Enriched Apoptosis Genes** |  |  |
| --- | --- | --- |
| Expressed in JM (not OC) | Expressed in OC (not JM) | Expressed in both JM & OC |
| Ank | Bgn | Clu |
| Atf3 | Birc3 | Dap |
| Birc | Casp4 | F2r |
| Cd44 | Cd14 | Hspb1 |
| Cdkn1a | Cd2 | Ifitm3 |
| Hmgb2 | Dcn | Lgals3 |
| Irf1 | Plcb2 | Mmp2 |
| Pea15a | Satb1 | Tgfb2 |
| Rela | Wee1 | Top2a |
| Rhob |  |  |
| Slc20a1 |  |  |
| Tnfrsf12a |  |  |

## Notes

### Competing Interest Statement

The authors have declared no competing interest.

### Summary of Updates

Additional Data / Figures have been added and an additional author has been added.

## REFERENCES

1. Rumballe BA, Georgas KM, Combes AN, et al. Nephron formation adopts a novel spatial topology at cessation of nephrogenesis. Dev Biol 2011; 360: 110–122.

2. Nosek TM. Essentials of Human Physiology, vol. Section 7/7ch03/7ch03p16. Gold Standard Multimedia Incorporated, 1998.

3. Ichii O, Yabuki A, Ojima T, et al. Rodent renal structure differs among species. J Vet Med Sci 2006; 68: 439–445.

4. Puelles VG, Hoy WE, Hughson MD, et al. Glomerular number and size variability and risk for kidney disease. Curr Opin Nephrol Hypertens 2011; 20: 7–15.

5. Jennette JC, D’Agati VD, Fogo AB, et al.: Heptinstall’s pathology of the kidney. In, Eighth edition. ed, Philadelphia, Wolters Kluwer,, 2024, p 1 online resource

6. Reilly R, Bulger R, Kriz W. Structural-functional relationships in the kidney. Diseases of the kidney and urinary tract 2007; 1: 2–53.

7. Denic A, Ricaurte L, Lopez CL, et al. Glomerular Volume and Glomerulosclerosis at Different Depths within the Human Kidney. J Am Soc Nephrol 2019; 30: 1471–1480.

8. Newbold KM, Sandison A, Howie AJ. Comparison of size of juxtamedullary and outer cortical glomeruli in normal adult kidney. Virchows Arch A Pathol Anat Histopathol 1992; 420: 127–129.

9. Samuel T, Hoy WE, Douglas-Denton R, et al. Determinants of glomerular volume in different cortical zones of the human kidney. J Am Soc Nephrol 2005; 16: 3102–3109.

10. Denic A, Mathew J, Lerman LO, et al. Single-Nephron Glomerular Filtration Rate in Healthy Adults. N Engl J Med 2017; 376: 2349–2357.

11. Bruns FJ, Alexander EA, Riley AL, et al. Superficial and juxtamedullary nephron function during saline loading in the dog. J Clin Invest 1974; 53: 971–979.

12. Bertram JF. Counting in the kidney. Kidney Int 2001; 59: 792–796.

13. Wiggins RC. The spectrum of podocytopathies: a unifying view of glomerular diseases. Kidney Int 2007; 71: 1205–1214.

14. Schneider RR, Eng DG, Kutz JN, et al. Compound effects of aging and experimental FSGS on glomerular epithelial cells. Aging (Albany NY) 2017; 9: 524–546.

15. Wright FS, Giebisch G. Glomerular filtration in single nephrons. Kidney Int 1972; 1: 201–209.

16. Pennell JP, Bourgoignie JJ. Adaptive changes of juxtamedullary glomerular filtration in the remnant kidney. Pflugers Arch 1981; 389: 131–135.

17. Ericson AC, Sjoquist M, Ulfendahl HR. Heterogeneity in regulation of glomerular function. Acta Physiol Scand 1982; 114: 203–209.

18. Horster M, Thurau K. Micropuncture studies on the filtration rate of single superficial and juxtamedullary glomeruli in the rat kidney. Pflugers Arch Gesamte Physiol Menschen Tiere 1968; 301: 162–181.

19. Roscioni SS, Heerspink HJ, de Zeeuw D. The effect of RAAS blockade on the progression of diabetic nephropathy. Nat Rev Nephrol 2014; 10: 77–87.

20. Rosenberg AZ, Kopp JB. Focal Segmental Glomerulosclerosis. Clin J Am Soc Nephrol 2017; 12: 502–517.

21. Haruhara K, Kanzaki G, Sasaki T, et al. Associations between nephron number and podometrics in human kidneys. Kidney Int 2022; 102: 1127–1135.

22. Romoli S, Angelotti ML, Antonelli G, et al. CXCL12 blockade preferentially regenerates lost podocytes in cortical nephrons by targeting an intrinsic podocyte-progenitor feedback mechanism. Kidney Int 2018; 94: 1111–1126.

23. Chen A, Sheu LF, Ho YS, et al. Experimental focal segmental glomerulosclerosis in mice. Nephron 1998; 78: 440–452.

24. Kastner S, Wilks MF, Gwinner W, et al. Metabolic heterogeneity of isolated cortical and juxtamedullary glomeruli in adriamycin nephrotoxicity. Ren Physiol Biochem 1991; 14: 48–54.

25. Sofue T, Kiyomoto H, Kobori H, et al. Early treatment with olmesartan prevents juxtamedullary glomerular podocyte injury and the onset of microalbuminuria in type 2 diabetic rats. Am J Hypertens 2012; 25: 604–611.

26. Iversen BM, Amann K, Kvam FI, et al. Increased glomerular capillary pressure and size mediate glomerulosclerosis in SHR juxtamedullary cortex. Am J Physiol 1998; 274: F365–373.

27. Edwards A, Kurtcuoglu V. Renal blood flow and oxygenation. Pflugers Arch 2022; 474: 759–770.

28. Chen J, Fleming JT. Juxtamedullary afferent and efferent arterioles constrict to renal nerve stimulation. Kidney Int 1993; 44: 684–691.

29. Chan GC, Eng DG, Miner JH, et al. Differential expression of parietal epithelial cell and podocyte extracellular matrix proteins in focal segmental glomerulosclerosis and diabetic nephropathy. Am J Physiol Renal Physiol 2019; 317: F1680–F1694.

30. Pippin JW, Sparks MA, Glenn ST, et al. Cells of renin lineage are progenitors of podocytes and parietal epithelial cells in experimental glomerular disease. Am J Pathol 2013; 183: 542–557.

31. Shigehara T, Zaragoza C, Kitiyakara C, et al. Inducible podocyte-specific gene expression in transgenic mice. J Am Soc Nephrol 2003; 14: 1998–2003.

32. Yamagami M, McFadden PW, Koethe SM, et al. Failure of T cell receptor-anti-CD3 monoclonal antibody interaction in T cells from marrow recipients to induce increases in intracellular ionized calcium. J Clin Invest 1990; 86: 1347–1351.

33. Marino S, Vooijs M, van Der Gulden H, et al. Induction of medulloblastomas in p53-null mutant mice by somatic inactivation of Rb in the external granular layer cells of the cerebellum. Genes Dev 2000; 14: 994–1004.

34. Wang Y, Eng DG, Kaverina NV, et al. Global transcriptomic changes occur in aged mouse podocytes. Kidney Int 2020; 98: 1160–1173.

35. Eng DG, Kaverina NV, Schneider RRS, et al. Detection of renin lineage cell transdifferentiation to podocytes in the kidney glomerulus with dual lineage tracing. Kidney Int 2018; 93: 1240–1246.

36. Lake BB, Menon R, Winfree S, et al. An atlas of healthy and injured cell states and niches in the human kidney. *bioRxiv* 2021: 2021.2007.2028.454201.

37. Zheng GX, Terry JM, Belgrader P, et al. Massively parallel digital transcriptional profiling of single cells. Nat Commun 2017; 8: 14049.

38. Wolf FA, Angerer P, Theis FJ. SCANPY: large-scale single-cell gene expression data analysis. Genome Biol 2018; 19: 15.

39. Wolock SL, Lopez R, Klein AM. Scrublet: Computational Identification of Cell Doublets in Single-Cell Transcriptomic Data. Cell Syst 2019; 8: 281–291 e289.

40. Lopez R, Regier J, Cole MB, et al. Deep generative modeling for single-cell transcriptomics. Nat Methods 2018; 15: 1053–1058.

41. Gayoso A, Lopez R, Xing G, et al. A Python library for probabilistic analysis of single-cell omics data. Nat Biotechnol 2022; 40: 163–166.

42. Traag VA, Waltman L, van Eck NJ. From Louvain to Leiden: guaranteeing well-connected communities. Sci Rep 2019; 9: 5233.

43. Li H, Dixon EE, Wu H, et al. Comprehensive single-cell transcriptional profiling defines shared and unique epithelial injury responses during kidney fibrosis. Cell Metab 2022; 34: 1977–1998 e1979.

44. Ritchie ME, Phipson B, Wu D, et al. limma powers differential expression analyses for RNA-sequencing and microarray studies. Nucleic Acids Res 2015; 43: e47.

45. Dill-McFarland KA, Mitchell K, Batchu S, et al. Kimma: flexible linear mixed effects modeling with kinship covariance for RNA-seq data. Bioinformatics 2023; 39.

46. Liberzon A, Birger C, Thorvaldsdottir H, et al. The Molecular Signatures Database (MSigDB) hallmark gene set collection. Cell Syst 2015; 1: 417–425.

47. Wang Y, Eng DE, Kaverina NV, et al. Global Transcriptomic Changes In Aged Mouse Podocytes. Kidney Int 2020; 95 (5): 1160–1173.

48. Pippin JW, Kaverina N, Wang Y, et al. Upregulated PD-1 signaling antagonizes glomerular health in aged kidneys and disease. J Clin Invest 2022; 132.

49. Pippin JW, Loretz CJ, Eng DG, et al. Isolation of Podocyte Cell Fractions From Mouse Kidney Using Magnetic Activated Cell Sorting (MACS). Bio Protoc 2025; 15: e5364.

50. Marshall CB, Krofft RD, Pippin JW, et al. CDK inhibitor p21 is prosurvival in adriamycin-induced podocyte injury, in vitro and in vivo. Am J Physiol Renal Physiol 2010; 298: F1140–1151.

51. Kaverina NV, Eng DG, Miner JH, et al. Parietal epithelial cell differentiation to a podocyte fate in the aged mouse kidney. Aging (Albany NY) 2020; 12: 17601–17624.

52. Kaverina N, Schweickart RA, Chan GC, et al. Inhibiting NLRP3 signaling in aging podocytes improves their life- and health-span. Aging (Albany NY) 2023; 15: 6658–6689.

53. Weidemann S, Bohle JL, Contreras H, et al. Napsin A Expression in Human Tumors and Normal Tissues. Pathol Oncol Res 2021; 27: 613099.

54. Veloso Pereira BM, Zeng Y, Maggiore JC, et al. Podocyte Injury at Young Age Causes Premature Senescence and Worsens Glomerular Aging. Am J Physiol Renal Physiol 2023.

55. Chen G, Cairelli MJ, Kilicoglu H, et al. Augmenting microarray data with literature-based knowledge to enhance gene regulatory network inference. PLoS Comput Biol 2014; 10: e1003666.

56. Fischer M. Census and evaluation of p53 target genes. Oncogene 2017; 36: 3943–3956.

57. Subramanian A, Tamayo P, Mootha VK, et al. Gene set enrichment analysis: a knowledge-based approach for interpreting genome-wide expression profiles. Proc Natl Acad Sci U S A 2005; 102: 15545–15550.

58. Padvitski T, Unger Avila P, Chen H, et al. Single-Cell Resolution of Cellular Damage Illuminates Disease Progression. bioRxiv 2025: 2025.2002.2020.639269.

59. Deleersnijder D, Venken T, Schepers R, et al. Single-Nucleus RNA-Sequencing Identifies a Differential Profibrotic Response in Parietal Epithelial Cells in Primary Versus Maladaptive Focal Segmental Glomerulosclerosis. Kidney Int Rep 2025; 10: 3255–3270.

60. Lennon R, Randles MJ, Humphries MJ. The importance of podocyte adhesion for a healthy glomerulus. Front Endocrinol (Lausanne) 2014; 5: 160.

61. Cina DP, Ketela T, Brown KR, et al. Forward genetic screen in human podocytes identifies diphthamide biosynthesis genes as regulators of adhesion. Am J Physiol Renal Physiol 2019; 317: F1593–F1604.

62. Maier JI, Rogg M, Helmstadter M, et al. EPB41L5 controls podocyte extracellular matrix assembly by adhesome-dependent force transmission. Cell Rep 2021; 34: 108883.

63. Rogg M, Maier JI, Dotzauer R, et al. SRGAP1 Controls Small Rho GTPases To Regulate Podocyte Foot Process Maintenance. J Am Soc Nephrol 2021; 32: 563–579.

64. Naylor RW, Watson E, Williamson S, et al. Basement membrane defects in CD151-associated glomerular disease. Pediatr Nephrol 2022; 37: 3105–3115.

65. Shankar PB, Nada R, Joshi K, et al. Podocin and beta dystroglycan expression to study podocyte-podocyte and basement membrane matrix connections in adult protienuric states. Diagn Pathol 2014; 9: 40.

66. McKinzie SR, Kaverina N, Schweickart RA, et al. Podocytes from hypertensive and obese mice acquire an inflammatory, senescent, and aged phenotype. Am J Physiol Renal Physiol 2024; 326: F644–F660.

67. Brinkkoetter PT, Bork T, Salou S, et al. Anaerobic Glycolysis Maintains the Glomerular Filtration Barrier Independent of Mitochondrial Metabolism and Dynamics. Cell Rep 2019; 27: 1551–1566 e1555.

68. Zhang J, Pippin JW, Krofft RD, et al. Podocyte Repopulation by Renal Progenitor Cells Following Glucocorticoids Treatment in Experimental FSGS. Am J Physiol Renal Physiol 2013; 304: F1375–1389.

69. Taniguchi Y, Pippin JW, Hagmann H, et al. Both cyclin I and p35 are required for maximal survival benefit of Cyclin-dependent kinase 5 in kidney podocytes. American Journal of Physiology- Renal Physiology 2012.

70. Marshall CB, Krofft RD, Blonski MJ, et al. Role of smooth muscle protein SM22alpha in glomerular epithelial cell injury. Am J Physiol Renal Physiol 2011; 300: F1026–1042.

71. Brinkkoetter PT, Wu JS, Ohse T, et al. p35, the non-cyclin activator of Cdk5, protects podocytes against apoptosis in vitro and in vivo. Kidney International 2010; 77: 690–699.

72. Brinkkoetter PT, Olivier P, Wu JS, et al. Cyclin I activates Cdk5 and regulates expression of Bcl-2 and Bcl-XL in postmitotic mouse cells. The Journal of clinical investigation 2009.

73. He J, Dijkstra KL, Bakker K, et al. Glomerular clusterin expression is increased in diabetic nephropathy and protects against oxidative stress-induced apoptosis in podocytes. Sci Rep 2020; 10: 14888.

74. Sanchez-Nino MD, Sanz AB, Sanchez-Lopez E, et al. HSP27/HSPB1 as an adaptive podocyte antiapoptotic protein activated by high glucose and angiotensin II. Lab Invest 2012; 92: 32–45.

75. Guo Y, Shen R, Yu L, et al. Roles of galectin–3 in the tumor microenvironment and tumor metabolism (Review). Oncol Rep 2020; 44: 1799–1809.

76. Shankland SJ, Freedman BS, Pippin JW. Can podocytes be regenerated in adults? Curr Opin Nephrol Hypertens 2017; 26: 154–164.

77. Kaverina NV, Eng DG, Schneider RR, et al. Partial podocyte replenishment in experimental FSGS derives from nonpodocyte sources. Am J Physiol Renal Physiol 2016; 310: F1397–1413.

78. Lasagni L, Angelotti ML, Ronconi E, et al. Podocyte Regeneration Driven by Renal Progenitors Determines Glomerular Disease Remission and Can Be Pharmacologically Enhanced. Stem Cell Reports 2015; 5: 248–263.

79. Padvitski T, Unger Avila P, Chen H, et al. Single-Cell Resolution of Cellular Damage Illuminates Disease Progression. bioRxiv 2025: 2025.2002.2020.639269.

80. Santo B, Fink EE, Krylova AE, et al. Exploring the utility of snRNA-seq in profiling human bladder tissue: A comprehensive comparison with scRNA-seq. iScience 2025; 28: 111628.

81. Wu H, Kirita Y, Donnelly EL, et al. Advantages of Single-Nucleus over Single-Cell RNA Sequencing of Adult Kidney: Rare Cell Types and Novel Cell States Revealed in Fibrosis. J Am Soc Nephrol 2019; 30: 23–32.

